# EXO1-mediated ssDNA gap expansion is essential for ATR activation and to maintain viability in BRCA1-deficient cells

**DOI:** 10.1101/2024.01.22.576613

**Authors:** Néstor García-Rodríguez, María del Carmen Domínguez-Pérez, Pablo Huertas

## Abstract

DNA replication faces challenges from DNA lesions originated from endogenous or exogenous sources of stress, leading to the accumulation of single-stranded DNA (ssDNA) that triggers the activation of the ATR checkpoint response. To complete genome replication in the presence of damaged DNA, cells employ DNA damage tolerance mechanisms that operate not only at stalled replication forks but also at ssDNA gaps originated by repriming of DNA synthesis downstream of lesions. Here, we demonstrate that human cells accumulate post-replicative ssDNA gaps following replicative stress induction. These gaps, initiated by PrimPol repriming and expanded by the long-range resection factors EXO1 and DNA2, constitute the principal origin of the ssDNA signal responsible for ATR activation upon replication stress, in contrast to stalled forks. Furthermore, we show that EXO1-deficient cells exhibit marked sensitivity to translesion synthesis inhibition, a distinctive characteristic of mutations in proteins essential for repairing ssDNA gaps via template switching, such as BRCA1/2. Strikingly, EXO1 loss results in synthetic lethality when combined with BRCA1 deficiency, but not BRCA2. Indeed, BRCA1-deficient cells become addicted to the overexpression of *EXO1 DNA2* or *BLM*. This dependence on long-range resection unveils a new vulnerability of BRCA1-mutant tumors, shedding light on potential therapeutic targets for these cancers.

## INTRODUCTION

The genetic information encoded by DNA is under constant threat from various internal and external sources of stress. One particularly vulnerable situation occurs during the process of DNA replication, where the replication machinery faces numerous obstacles that can lead to what is known as “replication stress” (1). Stalling of the replication forks results in the formation of stretches of single-stranded DNA (ssDNA), likely as a consequence of the uncoupling of helicase and polymerase activities or leading and lagging strand synthesis. The persistent accumulation of ssDNA, bound by Replication Protein A (RPA), triggers the activation of the replication stress response, primarily mediated by the kinase ataxia-telangiectasia and Rad3-related (ATR) (2). Once ATR is activated, it phosphorylates the effector kinase CHK1, which modulates the checkpoint response that ultimately leads to cell cycle arrest and DNA repair (3).

To allow the resumption of DNA replication in case of fork stalling, cells have evolved DNA damage tolerance (DDT) mechanisms. These mechanisms ensure the complete replication of the genome while preventing the deleterious consequences of prolonged fork stalling such as fork collapse and, ultimately, genome instability and/or cell death (4, 5). One of the DDT mechanisms is known as translesion DNA synthesis (TLS), which employs Y-family DNA polymerases (Polη, Polτ, Polκ and REV1). These specialized polymerases are able to bypass DNA lesions by inserting nucleotides opposite the damaged site, albeit in a potentially error-prone manner. Following this insertion step, DNA is extended by the B-family DNA polymerase Polζ, composed of REV3L, REV7, POLD2 and POLD3 (6, 7). Another mechanism of DDT is template switching (TS), an error-free recombinational process that uses the nascent DNA daughter strand on the sister chromatid as a temporary template to bypass the DNA lesion (5). Moreover, stalled replication forks can undergo a structural rearrangement known as replication fork reversal, where the two daughter strands anneal, leading to the formation of a four-way junction structure (5). Fork reversal serves to impede fork collapse while also triggering TS by enabling the blocked 3’end to use the complementary daughter strand for DNA extension beyond the point of stalling. As an alternative to TLS or TS, a third mechanism of DDT involves the re-initiation of DNA synthesis beyond the lesion through repriming. This mechanism leads to the formation of an unreplicated ssDNA gap behind the advancing fork, which subsequently need to be filled in a post-replicative manner through either TLS or TS, a process referred to as gap filling or post-replication repair (4, 8). In vertebrates, repriming is thought to be catalyzed by the Primase-Polymerase PrimPol (9–11). How cells choose among the different DTT pathways is currently under active investigation. In budding yeast, it has been recently shown that most of the ssDNA signal accumulating during replication upon methyl methanesulfonate (MMS) or ultraviolet (UV) damage arises at postreplicative regions, thus suggesting repriming as a predominant mechanism of DDT (12). Differently from budding yeast, replication fork reversal, detected by electron microscopy, was found to be a global response to replication challenges in mammalian cells (13). However, recent studies have confronted this model by revealing that the response to genotoxic stress varies depending on factors such as the nature or dosage of the replication challenge (4, 14). In this regard, the formation of ssDNA gaps through repriming has been observed in mammalian cells following the induction of a wide range of DNA lesions, including bulky adducts (15, 16) or DNA interstrand crosslinks (17, 18), and even upon other sources of replication stress that do not induce DNA damage, such as the depletion of nucleotide pools, inhibition of PARP1, or DNA secondary structures (19–21).

The coordination between checkpoint signaling and fork restart mechanisms is poorly understood. The extent of fork uncoupling can be limited by repriming events that can reduce the exposure of ssDNA at forks. This activity is expected to hinder the activation of the checkpoint, which requires a critical number of arrested forks (22, 23). In yeast, it has been observed that the ssDNA signal triggering the activation of the replication checkpoint upon replisome-stalling lesions primarily originates from the expansion of post-replicative gaps rather than directly form the fork (24). However, it is yet to be determined whether checkpoint signaling also arises from gaps in human cells. Replication gaps have emerged as a potential vulnerability in cancer cells (14, 19, 25). Intriguingly, deficiencies in the hereditary breast cancer genes, *BRCA1* and *BRCA2*, have been found to result in the accumulation of ssDNA gaps, attributed to defects in Okazaki fragment processing or gap repair mechanisms (14, 19, 26–29). More recently, we participated in a study that showed that the BRCA1/BARD1 ubiquitin ligase ubiquitinates PCNA to promote continuous DNA synthesis, avoiding the generation of ssDNA gaps (30). In addition, treatment with PARP inhibitors (PARPi), a common therapy for homologous recombination (HR)-deficient tumors, leads to the formation of ssDNA gaps in BRCA1-deficient cells, while the emergence of PARPi resistance appears to correlate with the suppression of these gaps (19, 31). Hence, targeting the formation and/or repair of ssDNA gaps could hold significant potential to maximize the efficacy of genotoxic chemotherapy.

Here, we provide evidence that ssDNA gaps readily emerge upon replication stress induced by the alkylating agent MMS or the replication inhibitor HU. These gaps are initiated by PrimPol-mediated repriming and subsequently expanded by the exonuclease EXO1, as well as the nuclease/helicase DNA2, thereby facilitating proper damage signaling through ATR activation. Of particular interest, we found that cells lacking EXO1, similar to BRCA1/2-deficient cells (26), exhibit an increased sensitivity to inhibition of the REV1-Polζ-dependent TLS. Remarkably, EXO1 is essential in BRCA1-deficient cells, but not in BRCA2-deficient cells. Thus, our findings open the possibility to target EXO1 as a therapeutic approach to treat BRCA1-deficient tumors.

## MATERIAL AND METHODS

### Cell lines and growth conditions

RPE1 h-TERT cells were grown in high-glucose DMEM/F-12 medium with L-glutamine (Sigma-Aldrich) supplemented with 10% fetal bovine serum (Sigma-Aldrich), 100 units/ml penicillin, and 100 μg/ml streptomycin (Sigma-Aldrich). RPE1 cells stably expressing YFP, YFP-EXO1 or YFP-*exo1*-*PIP* plasmids were grown in standard RPE1 medium supplemented with 0.5 mg/ml G418 (Gibco).

U2OS and HeLa cells were grown in high-glucose DMEM medium with L-glutamine (Sigma Aldrich) supplemented with 10% fetal bovine serum (Sigma-Aldrich), 100 units/ml penicillin, and 100 μg/ml streptomycin (Sigma-Aldrich).

MDA-MD-231, MDA-MB-436, DLD-1 and DLD-1 *BRCA2*^-/-^ cells were grown in RPMI-1640 without L-glutamine (Sigma-Aldrich), supplemented with L-Glutamine (Gibco), 10% fetal bovine serum (Sigma-Aldrich) 100 units/ml penicillin, and 100 μg/ml streptomycin (Sigma-Aldrich).

All cells were maintained at 37°C in 5% CO_2_. Cells were regularly tested for mycoplasma contamination.

### siRNAs, plasmids and transfections

siRNAduplexes were obtained from Sigma-Aldrich or Dharmacon (Supplementary Table S1) and were transfected using RNAiMax Lipofectamine Reagent Mix (Life Technologies), according to the manufactureŕs instructions.

YFP-EXO1 plasmid was a gift from Lene J. Rasmussen (University of Copenhagen) (32). YFP-*exo1-PIP* mutant plasmid was generated by mutating the previously published PIP box on EXO1 (33), using the QuickChange Lightening Site-Directed Mutagenesis kit (Agilent Technologies) according to the manufacturés guidelines. Plasmid pEYFP-N1 (Clontech) was used as control. Plasmid transfections were carried out using FuGENE 6 Transfection Reagent (Promega) according to the manufactureŕs protocol.

### Generation of EXO1 KO cell line

For the generation of EXO1 KO cell line, a predesigned guide RNA (gRNA) from Integrated DNA Technologies (IDT) targeting the sequence 5’-GCGTGGGATTGGATTAGCAA located in exon 8 of the EXO1 gene was used. RPE1 hTERT cells were transfected with a Cas9:gRNA ribonucleoprotein complex (IDT) according to the manufacturer’s guidelines. After 24 hours, transfected cells were seeded at low confluency to allow single-clone colony formation. Isolated colonies were tested for EXO1 expression by western blot. To test putative EXO1 KO clones, the CRISPR/Cas9 target site was amplified with specific primers hEXO1 CRISPR test AA down (5’ TCAAGCTCGGCTAGGAATGT) and hEXO1 CRISPR test AA up (5’ CTGCCGGGACTCAAAAAGT) and sequenced.

### Chromatin fractionation

Cells were harvested and resuspended in 180 µl of Cell fractionation buffer A (10 mM HEPES pH 7.5, 10 mM KCl, 1.5 mM MgCl_2_, 0.34 M sucrose, 10% glycerol, 1 mM DTT, complemented with 1x protease inhibitor cocktail (Roche)) + 0.1% Triton X-100, followed by incubation on ice for 15 min. At this point, a fraction of the volume was collected as whole cell extract (WCE), while the remaining volume was centrifuged at 1300g for 5 min at 4°C to precipitate the nuclei. The pellet was washed with buffer A and gently resuspended in 180 µl of Cell fractionation buffer B (20 mM HEPES pH 7.5, 3 mM EDTA, 10% glycerol, 125 mM potassium acetate, 1.5 mM MgCl_2_, 1 mM DTT, 0.5% Triton X-100, complemented with 1x protease inhibitor cocktail) followed by incubation on ice for 30 min and centrifugation at 1700g for 5 min at 4°C to precipitate the chromatin. The pellet was washed once with buffer B, resuspended in 1x Laemmli sample buffer, and boiled at 100°C for 5 min (Chromatin sample). WCE and chromatin samples were subjected to western blotting analysis.

### Western blotting

Cells were collected by scrapping in Laemmli buffer 2x (4% SDS, 20% glycerol, 125 mM Tris-HCl, pH 6.8) and boiled for 5 min at 100°C. 4x Laemmli sample buffer with β-mercaptoethanol was added to the lysates, followed by denaturation at 100°C for 5 min. Proteins were then resolved by SDS-PAGE and transferred to an Amersham Protran nitrocellulose membrane (Cytvia Life Sciences). Following incubation for 1h in blocking buffer (3-5% milk in TBS + 0.1% Tween-20), membranes were incubated with the primary antibodies diluted in blocking buffer overnight at 4°C. Anti-CHK1-pS345 (Cell Signaling #2348), anti-CHK1 (Santa Cruz #8408) anti-EXO1 (GeneTex #109891), anti-β-Actin (Abcam #8227), anti-α-Tubulin (Sigma #T9026), anti-RPA2-pS4/pS8 (Bethyl #A300-245A), anti-RPA2 (Abcam #2175), anti-KAP1-pS824 (Bethyl #A300-767A), anti-γ-H2AX (Cell Signaling #2577), anti-MRE11 (Novus Biological #NB100-142), anti-PrimPol (11), anti-H3 (Abcam #1791), anti-BRCA1 (Santa Cruz #6954) and anti-BRCA2 (Millipore #OP95) primary antibodies were used. Membranes were then incubated with the appropriate infra-red dyed (LI-COR) or HRP-labelled secondary antibodies and scanned in an Odyssey Infra-red Imaging System (Li-COR) or a Chemidoc MP imaging system (BioRad). Images were analyzed and quantified with ImageStudio software (LI-COR).

### S1 nuclease assay

Cells were pulse-labeled with 20 µM IdU (20 min), washed twice with PBS and pulse-labeled with 200 µM CldU in the absence or presence of 1 mM MMS for 1h. Cells were then washed twice with PBS and permeabilized with CSK buffer (100 mM NaCl, 10 mM MOPS pH 7, 3 mM MgCl_2_, 300 mM Sucrose and 0.5% Triton X-100 in water) for 8 min at RT. Permeabilized cells were treated with S1 nuclease buffer (30 mM Sodium acetate pH 4.6, 10 mM Zinc acetate, 5% glycerol, 50 mM NaCl in water) with or without 20 U/ml S1 nuclease (Invitrogen, 18001-016) for 30 min at 37°C. Cells were then scrapped in PBS + 0.1% BSA, pelleted and resuspended in PBS + 0.1% BSA at a final concentration of 1-2×10^3^ cells/µl. 2.5 µl of cell suspension were spotted on a positively charged slide and lysed with 7.5 µl of spreading buffer (200 mM Tris-HCl pH 7.5, 50 mM EDTA, 0.5% SDS). After 8 min, slides were tilted at 45 degrees to allow the DNA to spread. Slides were then air-dried, fixed with ice-cold methanol/acetic acid (3:1) for 5 mins, air-dried and stored at 4°C. Slides were rehydrated with PBS, denatured with 2.5 M HCl for 1h, washed with PBS twice, and blocked with blocking buffer (3% BSA, 0.1% Triton X-100 in PBS) for 40 min. Next, slides were incubated with primary antibody mix of mouse anti-BrdU which recognizes IdU (Becton Dickinson #347580, 1:250) and rat anti-BrdU which recognizes CldU (Abcam #6326, 1:250) diluted in blocking buffer for 2.5h at RT in a dark humid chamber. Slides were washed 3 times with PBS for 5 min each and incubated with secondary antibodies anti-mouse Alexa fluor 594 and anti-rat Alexa fluor 488 (1:250, Invitrogen #A11032 and #A11006, respectively) in blocking buffer for 1h at RT in a dark humid chamber. After washing 3 times with PBS and air-drying, slides were mounted with Prolong gold antifade reagent (Invitrogen, P36930) and stored at 4°C until imaging. Images were acquired using a AF6000 Leica Fluorescence microscope equipped with a HCX PL APO 63x (NA =1.4) oil objective. Fibers were measured using the segmented line tool on ImageJ FIJI software (https://fiji.sc).

### Cell viability assays

Cells were seeded in 6-well plates at appropriate dilutions in triplicates. After 7-14 days, colonies were stained with a solution containing 0.5% crystal violet and 20% ethanol, followed by several washes with water. Plates were scanned with an Epson Perfection 4490 scanner and the area of the wells covered with cells was calculated using the ImageJ plugin ColonyArea (34). For viability assays upon drug treatment, 24h after seeding, the cells were treated with DMSO as control, or with the chemical agent described in the text at the indicated concentrations. After 7-14 days, colonies were stained, scanned, and quantified as indicated previously. For the clonogenic survival shown in Figure 5B, the number of colonies were counted manually. Data are representative of at least three independent experiments.

### Immunofluorescence

For RPA signal visualization, cells were seeded on coverslips. After 1 h of MMS treatment (2mM), cells were treated with pre-extraction buffer (25 mM Tris-HCl, pH 7.5, 50 mM NaCl, 1 mM EDTA, 3 mM MgCl_2_, 300 mM sucrose and 0.2% Triton X-100) for 5 min on ice to pre-extract soluble proteins. Cells were then fixed with 4% paraformaldehyde (w/v) in PBS for 20 min.

For 53BP1 visualization, cells were seeded on coverslips. After 1 h upon addition of the indicated concentrations of the genotoxic agents, cells were fixed with ice-cold methanol for 10 min, followed by incubation with ice-cold acetone for 30 s.

In both cases, after two washes with PBS, cells were blocked for 1h with 5% FBS in PBS, co-stained with anti-RPA (Abcam #21752) or anti-53BP1 (Novus Biologicals #NB100-304) primary antibodies diluted in blocking buffer for 2 h at room temperature, washed again with PBS and then co-immunostained with the secondary antibodies anti-mouse Alexa Fluor 594 (Invitrogen #A11032) for RPA or anti-rabbit Alexa Fluor 488 (Invitrogen #A11034) for 53BP1 in blocking buffer for 1 h at room temperature. Coverslips were then mounted into glass slides using Vectashield mounting medium with DAPI (Vector laboratories). Images were acquired using a AF6000 Leica Fluorescence microscope equipped with a HCX PL APO 63x (NA =1.4) oil objective. RPA signal intensities were quantified using ImageJ FIJI software.

### Flow cytometry-based immunofluorescence

Cells were collected by trypsinization and treated with 4% formaldehyde (w/v) for 15 min at room temperature. After washing with PBS, cells were pelleted by centrifugation, resuspended in ice-cold methanol, and incubated for 10 min on ice. Cells were then washed with PBS and incubated with anti-CHK1-pS345 in antibody dilution buffer (0.5% BSA in PBS) for 1.5 h. Following a wash with PBS, cells were co-immunostained with the secondary antibody anti-rabbit Alexa Fluor 488 (Invitrogen #A11034) in antibody dilution buffer for 45 min at room temperature, washed with PBS and incubated with 250 μg/ml RNase A (Sigma) and 1 μg/ml DAPI (Sigma) overnight. Samples were analyzed with a LSRFortessa X-20 machine (BD Biosciences). The histograms shown in Figure S2F were generated by gating the cell population in S-phase based on the DNA content.

### Gene expression correlation and survival analysis

Correlations between the expressions of *BRCA1/2*, *EXO1, DNA2, BLM* and *MRE11* and survival data from the TCGA Breast Cancer (BRCA) dataset were obtained using the UCSC Xena platform (35). Heatmaps and Kaplan-Meier survival analyses were generated using PRISM software (Graphpad Software). The comparison of *EXO1* expression levels between tumor and normal samples in TCGA Breast cancer and GTEx datasets was performed using the GEPIA platform (36).

### Cell cycle analysis

Cells were fixed with cold 70% ethanol overnight, incubated with 250 μg/ml RNase A (Sigma) and 10 μg/ml propidium iodide (Fluka) in PBS at 37°C for 30 min and analyzed with a FACSCalibur or a LSRFortessa X-20 machines (BD Biosciences).

### Statistical analysis

Statistical significance was determined with the test indicated in the corresponding figure legend using PRISM software (Graphpad Software). Statistically significant differences were labeled with one, two, three or four asterisks if p<0.05, p<0.01, p<0.001 or p<0.0001, respectively. Specific replicate numbers (n) for each experiment can be found in the corresponding figure legends.

## RESULTS

### Unreplicated ssDNA gaps are formed in response to replication stress

The equilibrium between different DDT pathways is still poorly understood. In several studies, it has been demonstrated that PrimPol repriming is selectively activated upon cisplatin or low doses of HU, under specific conditions such as impaired fork reversal, PrimPol overexpression or BRCA deficiency (17, 20, 25, 26, 37). However, the relevance of repriming in WT cells is less understood. We then decided to use non-transformed RPE1-hTERT cells (referred to as WT in this manuscript) to investigate whether repriming plays a role in the DNA replication stress response triggered by the alkylating agent methyl methanesulfonate (MMS). First, we exposed WT cells to MMS for 1 hour, and examined the recruitment of PrimPol to chromatin. We also exposed the cells to the replication inhibitor hydroxyurea (HU), which has been shown previously to increase the amount of PrimPol on chromatin (11). Notably, both MMS and HU resulted in the accumulation of PrimPol protein on the chromatin fraction (Figure 1A).

**Figure 1.**
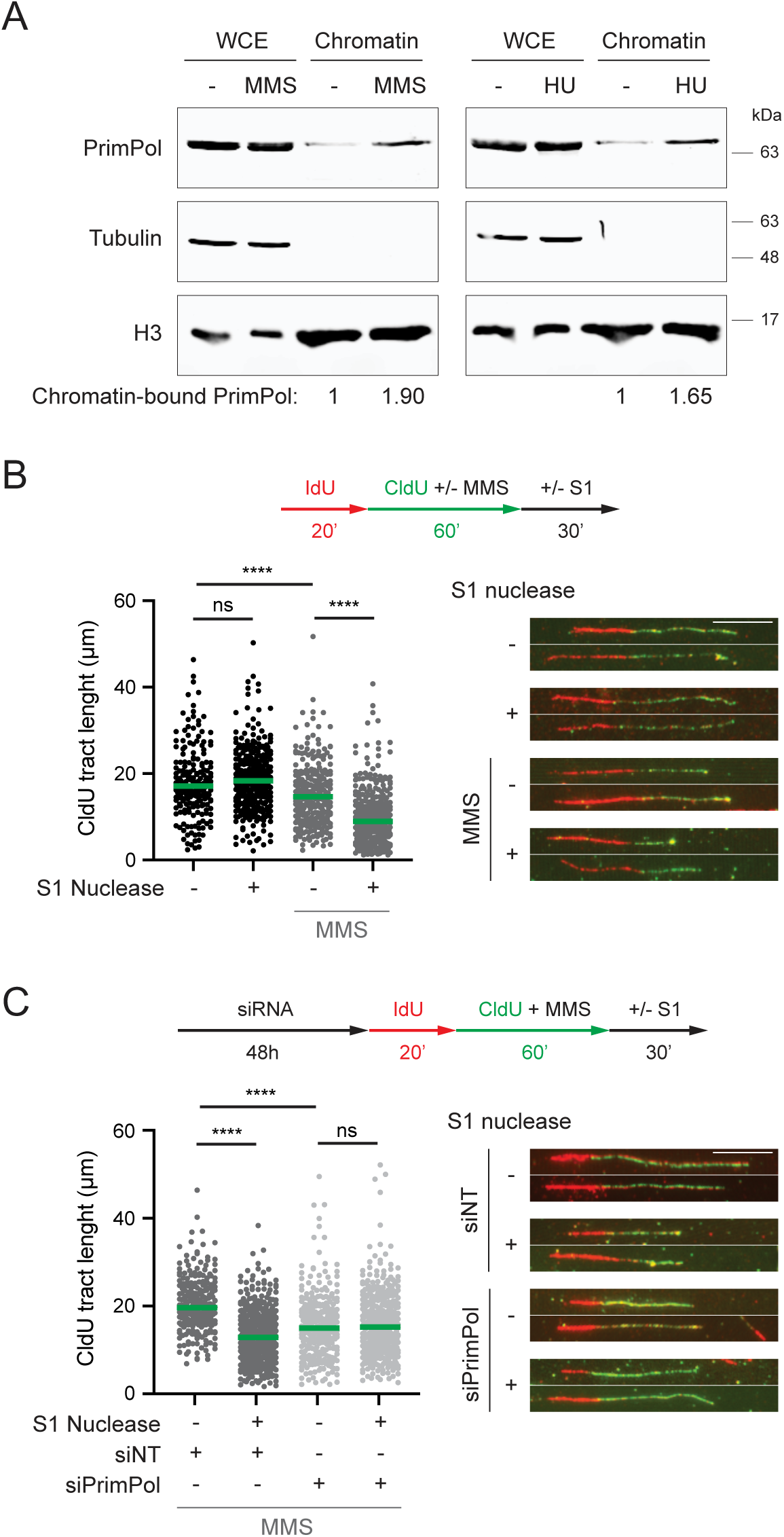
Unreplicated ssDNA gaps are formed in response to replication stress. **A.** Immunoblots show the levels of the indicated proteins in whole cell extracts (WCE) and chromatin-enriched fractions upon treatment with MMS (left, 2 mM) or HU (right, 2 mM) for 1h. Quantification of chromatin-bound PrimPol relative to histone H3 is shown on the bottom. **B** and **C**. Top: Scheme of the IdU/CldU pulse-labelling protocol, followed by S1 nuclease treatment. Bottom: CldU tact lengths in the indicated experimental conditions with or without S1 nuclease treatment. Each dot represents one fiber and the green bar represents the median. At least 200 fibers were measured from two biological experiments (n=2). p values were calculated using the Mann-Whitney test (ns: p>0.05; ****: p<0.0001). Representative images of DNA fibers are shown. Scale bar, 10 μM.

To precisely assess whether ssDNA gaps were generated in response to MMS, we used the S1 nuclease DNA fiber method (38). In this assay, after consecutive pulse-labelling with two thymidine analogs, IdU and CldU, cells are permeabilized and incubated with the S1 endonuclease that specifically cleaves ssDNA. This treatment shortens the DNA tracts if gaps are present in nascent DNA. Control samples showed that the CldU tract length remained unchanged upon S1 digestion. However, when MMS was added during the CldU labelling, the tracts became sensitive to S1 nuclease (Figure 1B), thereby indicating the formation of gaps within the DNA strands. Accordingly, depletion of PrimPol prevented the CldU tracts from shortening after S1 treatment (Figure 1C and S1A). Notably, in the absence of S1 nuclease treatment, CldU tracts were shorter in PrimPol-depleted cells compared to control cells (Figure 1C), suggesting a role of PrimPol during replication through MMS-induced lesions. In line with this observation, loss of PrimPol sensitized human RPE1 cells to MMS exposure (Figure S1B), consistent with previous findings in avian cells (39). Taken together, these data indicate that ssDNA gaps are readily generated upon MMS treatment through repriming by PrimPol. Interestingly, HU treatment has also been demonstrated to result in accumulation of ssDNA gaps in budding yeast, as well as in BRCA1/2-deficient human cells (25, 40). Therefore, our findings, along with those of others, suggest that the formation of ssDNA gaps via PrimPol repriming activity is a common response to any form of replication stress even in wildtype human cells.

### EXO1 activity is required for robust ATR activation

In budding yeast, the replication stress response elicited by polymerase-blocking lesions primarily arises not from the fork, but rather from daughter-strand gaps, left behind the replication forks. These ssDNA gaps need to be enlarged by various factors, including the exonuclease Exo1, in order to generate an adequate checkpoint signal (24). Based on our observations that ssDNA gaps are generated in human cells in response to replication stress, we wonder whether EXO1-mediated gap expansion may also be a pre-requisite for producing a sufficient checkpoint signaling. To address this question, we generated a CRISPR/Cas9-mediated EXO1 KO cell line in RPE-1-hTERT cells (referred to as *exo1* in this manuscript). We initially monitored the accumulation of ssDNA in both WT and *exo1* cells, using as a readout the signal intensity of the ssDNA-binding protein RPA within the nucleus, detected by immunofluorescence (Figure 2A). Notably, RPA mean intensity in RPA-positive cells, which correspond to those in S-phase, was highly increased in the WT after MMS treatment. However, in cells lacking EXO1, such increased in RPA intensity was significantly lower. Thus, this data suggests that EXO1 activity promotes the accumulation of ssDNA in the presence of damaged DNA.

**Figure 2.**
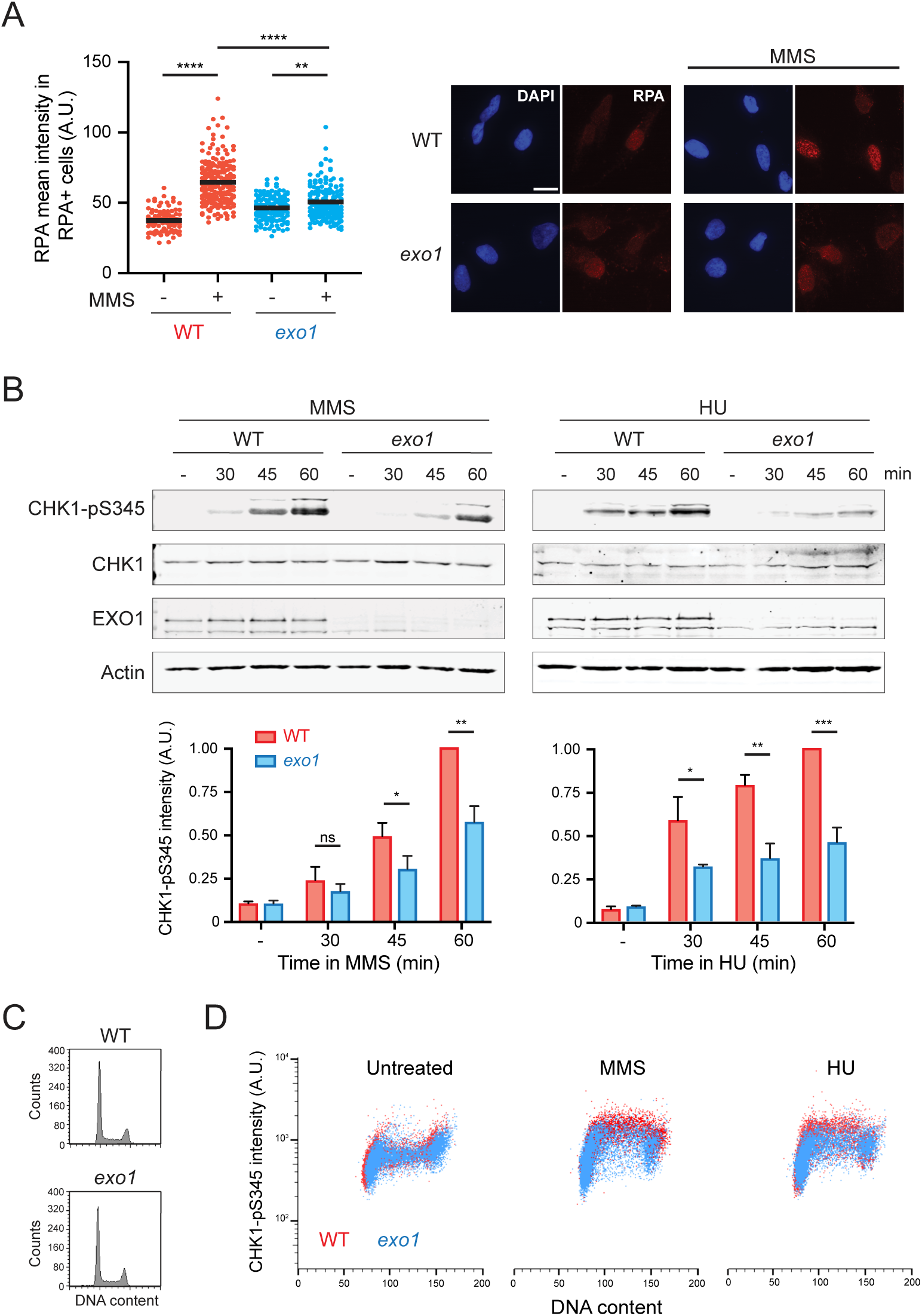
EXO1 function is required for ssDNA accumulation and, consequently, robust ATR checkpoint activation upon replicative stress. **A**. Quantification of the RPA mean signal intensity in the fraction of cells positive for RPA in WT and *exo1* cells treated with or without MMS (2 mM) for 1h. Each dot represents the RPA mean intensity inside the nucleus and the black bar represents the mean. Right: representative images. Scale bar, 20 μM. **B.** Western blots for monitoring CHK1-S345 phosphorylation in WT and *exo1* cells upon addition of MMS (left, 2 mM) or HU (right, 2 mM). Actin served as loading control. Quantifications of CHK1-pS345 intensities, normalized to actin, are shown below. Bars represent the means ± SD of independent biological replicates (n=3). p-values were calculated using the Student’s *t*-test (ns: p>0.05; *: p<0.05; **: p<0.01; ***: p<0.001). **C.** Cell cycle profiles of WT and *exo1* cells. **D.** Quantifications of CHK1-pS345 signal intensities in WT (red) and *exo1* (blue) cells by flow cytometry-based immunofluorescence, left untreated or treated with MMS (2 mM) or HU (2 mM). Each dot represents a single cell. 10, 000 cells are displayed per condition.

Next, we evaluated whether a reduction in the overall accumulation of ssDNA could potentially have an impact on the activation of the ATR checkpoint. As a readout for ATR activation, we analyzed the phosphorylation of CHK1 at Ser^345^ at very early timepoints upon addition of MMS, HU or 4-Nitroquinoline 1-oxide (4-NQO), an ultraviolet radiation-mimetic compound (Figure 2B and S2A). Indeed, the phosphorylation of CHK1 was significantly delayed in *exo1* when compared to wild-type cells. However, the observed difference was no longer evident after extending the treatment for up to three hours, during which the signal reached saturation (Figure S2B). This indicates that the requirement of EXO1 for the full activation of ATR depends on the extent of the damage. Additionally, the defect in ATR activation was further validated in a second KO clone of EXO1 in RPE-1 cells, as well as in the cancer cell lines U2OS and HeLa by siRNA-mediated depletion of EXO1 (Figure S2C-E). Although a decrease in the level of CHK1 phosphorylation could be explained by a reduction in the number of cells in S-phase, no significant changes were observed in the cell cycle profiles of EXO1-lacking cells compared to control (Figure 2C and Figure S2C-E). To completely rule out this possibility, we monitored the intensity of CHK1 phosphorylation in single cells using a flow cytometry-based immunofluorescence assay (Figure 2D and Figure S2F). After subjecting WT cells to HU or MMS treatments, we found a noticeable increase in CHK1-pS345 intensity from the background levels shown in untreated cells. This increase was particularly evident in the fraction of cells in S-phase. However, in *exo1* cells, this increase was relatively lower, which further support the idea that EXO1 is required for timely and robust ATR activation upon induction of replication stress in human cells.

EXO1 is involved in long-range resection during the homologous recombination pathway for double-strand break (DSB) repair, thus generating ssDNA from the DNA ends that activates the ATR checkpoint (41). Therefore, the observed delay in ATR activation in *exo1* cells could be attributed to an impairment in DNA-end resection following the potential formation of DSBs by the treatments with the different genotoxic agents. We then set to look for evidence of DSB formation under the moderate damage conditions used in this study. First, we conducted western blot experiments to examine the phosphorylation RPA2 at S4/S8 and KAP1 at S824 (Figure S3A). These phosphorylation events serve as indicators of ATM/DNA-PK activation and are commonly used as markers for DSBs (13). The topoisomerase I inhibitor camptothecin (CPT) was used as positive control for DSBs formation (42). Treatments for up to 1h with the selected doses of MMS, HU and 4-NQO did not reveal any significant increase in the phosphorylation levels of RPA2. However, a slight increase in phospho-KAP1 was detected after 1h, specially upon MMS, albeit considerably lower than the substantial increase observed upon treatment with CPT (Figure S3A). MMS and HU have been already shown to induce phosphorylation of KAP1 without detectable DSB formation and RPA2 S4/S8 phosphorylation, supporting the notion of DSB-independent activation of ATM by replication stress (13). Intriguingly, ATR activation, as detected by phosphorylation of CHK1, was similar following all the treatments, thus supporting the idea that ATR was activated by different structures (Figure S3A). Second, we analyzed the formation of 53BP1 foci, as a marker for the non-homologous end-joining (NHEJ) pathway for DSB repair (Figure S3B). In agreement with the western blot data, 53BP1 foci were detected 1h after the addition of CPT but not MMS, HU or 4-NQO. Collectively, these findings suggest that EXO1 facilitates ATR activation through the processing of structures other than DSBs, likely involving post-replicative ssDNA gaps.

### EXO1 binding to PCNA is required for robust ATR activation

EXO1 has been shown to interact with PCNA through a PCNA-interacting Protein region (PIP-Box)-like sequence located at the C-terminus of the protein (32). This interaction has been shown to promote EXO1 damage recruitment and processivity during DSB repair (33). We then sought to elucidate if the interaction of EXO1 with PCNA was required for ATR checkpoint activation upon induction of replicative stress. To do so, we generated stable cell lines expressing an empty vector, a YFP-tagged wild-type version of EXO1 gene (YFP-EXO1) or a YFP-tagged PIP box mutant of EXO1 (YFP-*exo1-PIP*) and monitored the activation of ATR upon MMS treatment by the phosphorylation of CHK1 (Figure 3). Accordingly, when compared to the WT, ATR activation was delayed in *exo1* cells expressing an empty vector, and this defect was nearly completely rescued in *exo1* cells expressing the wild-type EXO1 gene (Figure 3A-B). However, the PIP-box mutant of EXO1 failed to rescue this delay (Figure 3C), thus indicating that an interaction of EXO1 with PCNA is required for efficient ATR activation.

**Figure 3.**
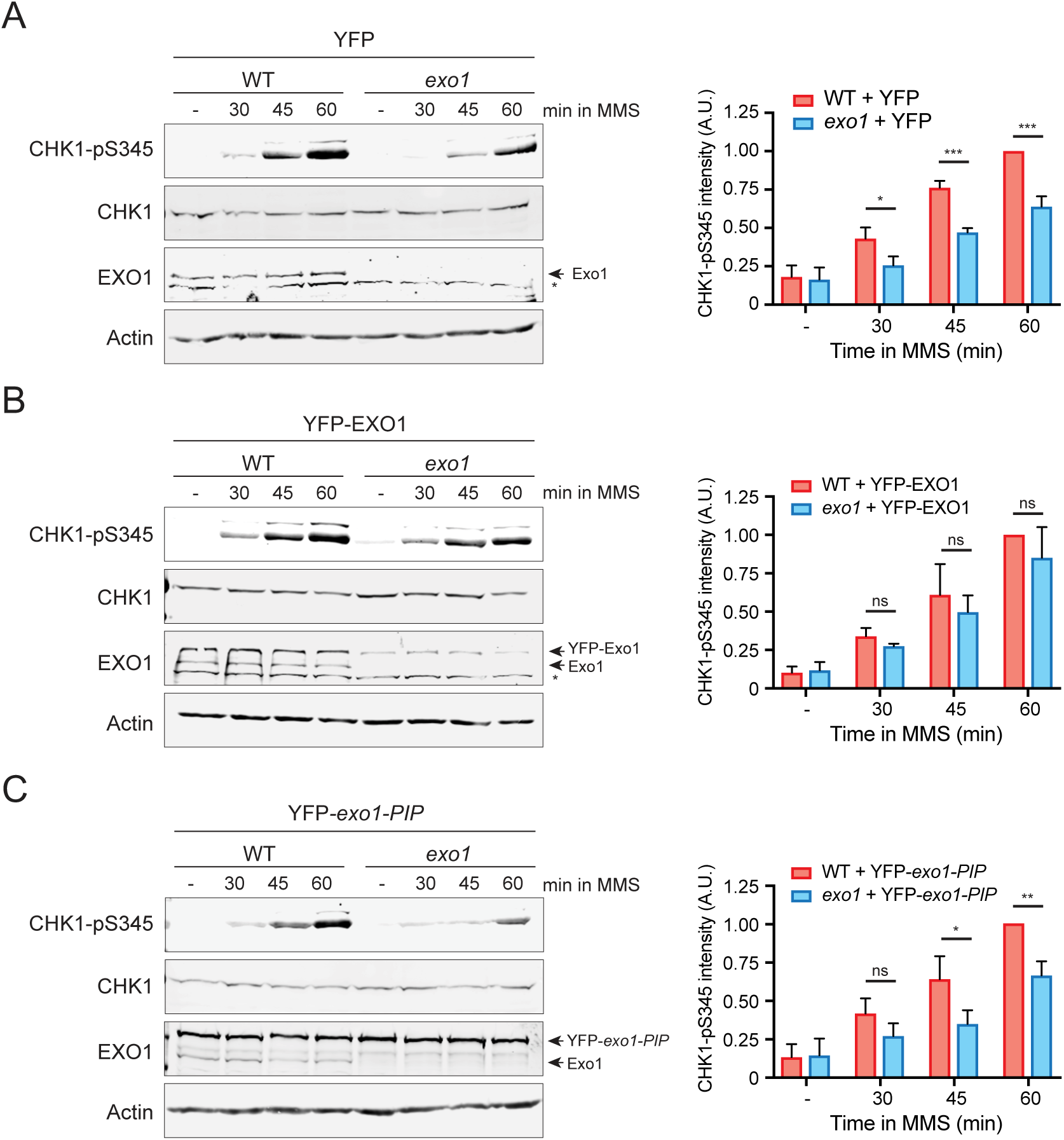
Interaction of EXO1 with PCNA is required for robust ATR checkpoint activation upon replicative stress. Western blots for monitoring CHK1-S345 phosphorylation in WT and *exo1* cells stably expressing an empty vector (**A**), a YFP-tagged EXO1 (**B**) and a YFP-tagged *exo1* PIP-box mutant (**C**), upon addition of MMS (2 mM). Actin served as loading control. Quantifications of CHK1-pS345 intensities, normalized to actin, are shown on the right. Bars represent the means ± SD of independent biological replicates (n=3). p-values were calculated using the Student’s *t*-test (ns: p>0.05; *: p<0.05; **: p<0.01; ***: p<0.001).

### DNA2, but not MRE11, activity is required for ATR activation

Next, we investigated whether the complete activation of ATR requires the involvement of other nucleases in addition to EXO1. First, we examined the role of MRE11, a component of the MRE11-RAD50-NBS1 (MRN) complex responsible for recognizing DSBs and initiating DNA-end resection (41, 43). Interestingly, MRE11 has also been implicated in the resection of ssDNA gaps, thereby facilitating RAD51 loading and subsequent homologous recombination (HR)-dependent repair of the gaps (16, 44). However, the depletion of MRE11 had no discernable impact on either the phosphorylation levels of CHK1 following MMS treatment or the cell cycle profile, when compared to the control samples (Figure S4A and S4B). Additionally, the inhibition of the exonuclease activity of MRE11 by mirin did not produce any noticeable impact on phospho-CHK1 levels following MMS exposure (Figure S4C).

Then, we focused on DNA2, a nuclease/helicase involved in long-range resection during the homologous recombination pathway for DSB repair (41, 43). To avoid the toxicity associated with siRNA-mediated depletion of DNA2, we used the specific DNA2 inhibitor C5 (45). Interestingly, the phosphorylation of CHK1 was significantly delayed upon MMS or HU treatment when DNA2 was inhibited (Figure 4A). Notably, the cell cycle profile of DNA2-inhibited cells remained similar to that of the control cells (Figure 4B), thereby ruling out any cell cycle contribution to the observed phenotype. Taken together, these data indicate that both EXO1 and DNA2 play a pivotal role as the primary nucleases involved in the generation of the ssDNA signal that activates the ATR checkpoint upon replication stress, whereas the impact of MRE11 appears to be more restricted.

**Figure 4.**
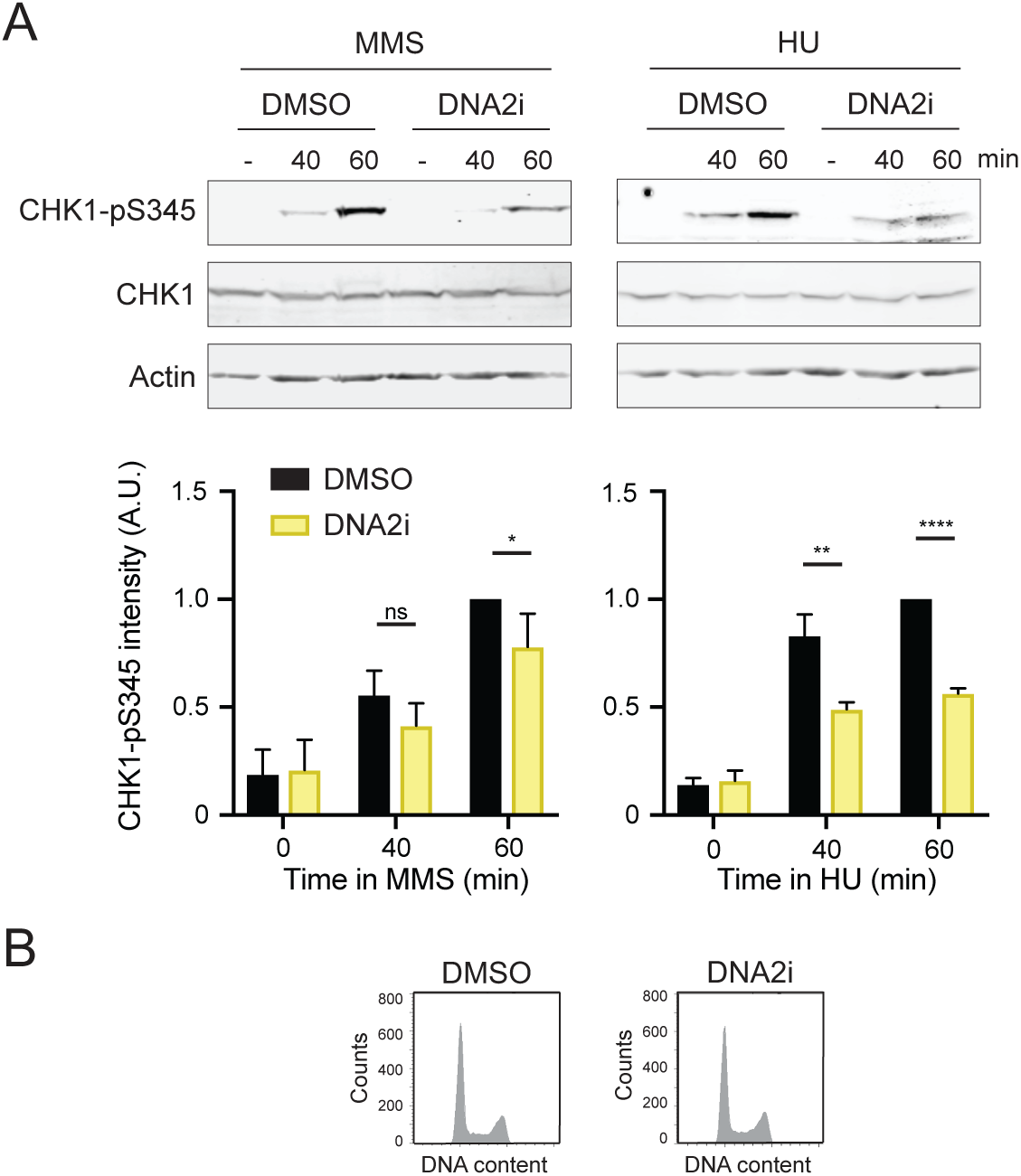
DNA2 activity is required for ATR checkpoint activation upon replication stress. **A**. Western blots for monitoring CHK1-S345 phosphorylation in WT (RPE1) cells treated with DMSO or DNA2 inhibitor C5 (20 μM) for 5h before the addition of MMS (2 mM) or HU (2 mM). Bottom panels: Quantification of CHK1-pS345 intensities, normalized to the loading control. Bars represent the means ± SD of independent biological replicates (n=4 for MMS, n=3 for HU). p-values were calculated using the Student’s *t*-test (ns: p>0.05; *: p<0.05; **: p<0.01; ****: p<0.0001). **B.** Cell cycle profiles in control (DMSO) or DNA2i-treated cells.

### EXO1 expands ssDNA gaps behind the replication forks

Our observations suggest that ssDNA gaps arising behind the replication forks in cells exposed to replication stress are expanded by EXO1 (and DNA2) nuclease, thereby contributing to the full activation of the ATR checkpoint in human cells. In order to test this hypothesis, we monitored the formation of ssDNA gaps in *exo1* cells by the S1 nuclease DNA fiber assay. We reasoned that if EXO1 expands ssDNA gaps, its absence would result in an increased probability that gaps remain undigested during S1 nuclease treatment. Indeed, while DNA tracts undergo shortening in WT cells treated with MMS when subjected to S1 nuclease digestion, they became insensitive to S1 nuclease in *exo1* cells (Figure 1B and 5A).

**Figure 5.**
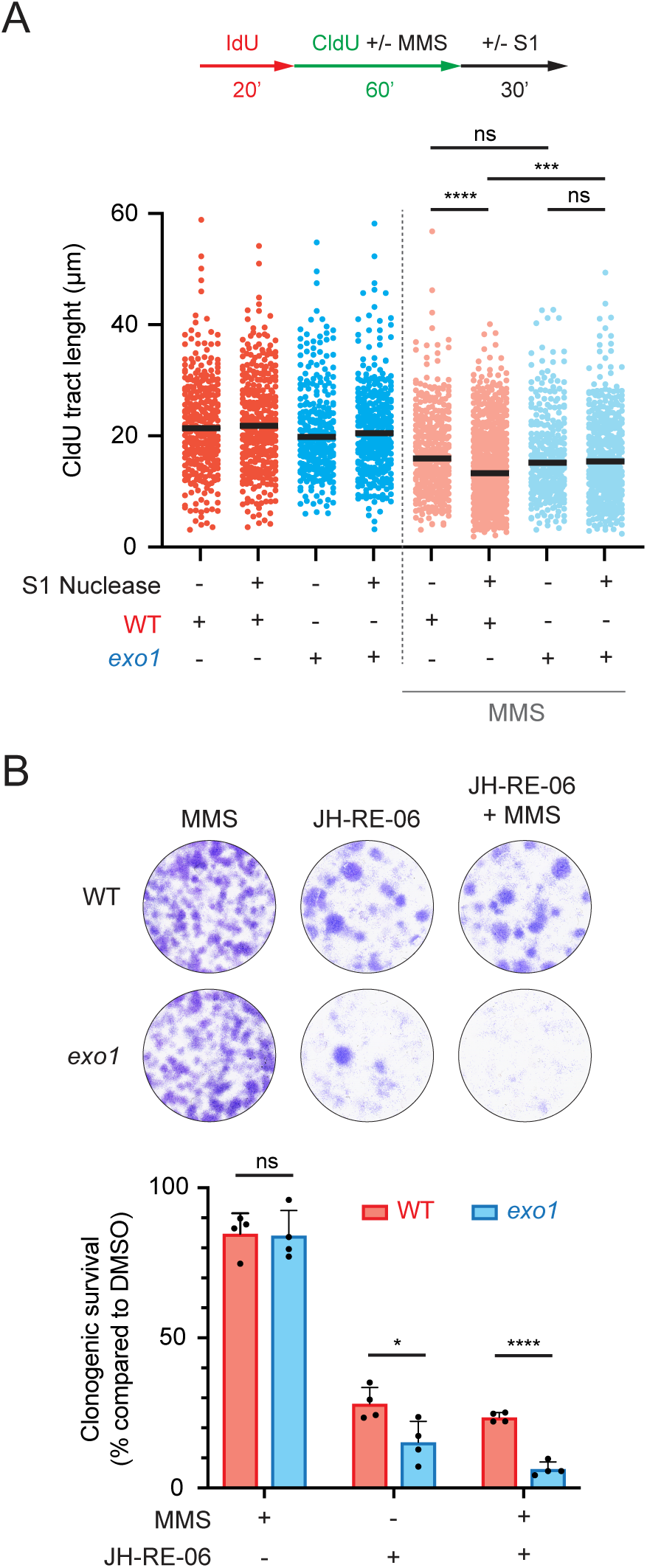
EXO1 facilitates the expansion of ssDNA gaps and promotes cell survival upon treatment with the REV1-Polζ inhibitor JH-RE-06. **A**. Top: Scheme of the IdU/CldU pulse-labelling protocol, followed by S1 nuclease treatment. Bottom: CldU tact lengths in the indicated experimental conditions with or without S1 nuclease treatment. Each dot represents one fiber and the green bar represents the median. At least 275 fibers were measured from two biological experiments (n=3). p values were calculated using the Mann-Whitney test (ns: p>0.05; ***: p<0.001; ****: p<0.0001). **B**. Clonogenic survival of WT and *exo1* cells treated with MMS (0.05 mM) and/or JH-RE-06 (0.25 μM). Top: Representative images. Bottom: Quantification of the clonogenic survival as a percentage of the number of colonies formed relative to control DMSO-treated cells. Bars represent the means ± SD of independent biological replicates (n=3). Values of individual experiments are indicated as dots. p values were calculated using the Student’s *t*-test (ns: p>0.05; *: p<0.05; ****: p<0.0001).

The expansion of ssDNA gaps has been proposed to be a prerequisite for successful template switching pathway of DNA damage tolerance (12, 16, 24, 46–48). When TS is impaired, cells tend to rely more on translesion synthesis for repairing ssDNA gaps arising during DNA replication. This is the case for HR-deficient cells, such as *BRCA1/2* mutant cells, which accumulate ssDNA gaps during unperturbed DNA replication and exhibit increased toxicity to the inhibition of the REV1-Polζ-dependent TLS (26, 30). We then tested the sensitivity of *exo1* cells to the REV1-Polζ inhibitor JH-RE-06 (Figure 5B). In agreement with a role of EXO1 in TS, *exo1* cells exhibited a considerably higher sensitivity to JH-RE-06 treatment compared to WT cells, resulting in a reduction of 50% in clonogenic survival. Interestingly, the sensitivity of *exo1* cells to JH-RE.06 was further enhanced when replicative stress was induced through chronic MMS treatment, leading to a further reduction in clonogenic survival when compared to WT cells (Figure 5B). Hence, these findings suggest that EXO1 plays a critical role in expanding ssDNA gaps generated during replication stress, a process essential for the subsequent repair by TS. Thus, a deficiency in TS renders *exo1* cells more reliant on TLS to deal with replication stress.

### EXO1 function is essential for BRCA1-deficient cells

Given that BRCA1/2 proteins promote the repair of ssDNA gaps through the TS pathway, and our data indicate that EXO1-mediated gap extension is crucial for TS, we set to investigate the potential repercussions of EXO1 loss in *BRCA1/2* mutant cells. Strikingly, depletion of EXO1 significantly reduced the viability of a BRCA1-deficient breast cancer cell line (MDA-MB-436) but had no effect on a BRCA1-proficient breast cancer cell line (MDA-MB-231) (Figure 6A and S5A). This reduction in viability was specific to BRCA1, as EXO1 depletion did not impact the survival of a BRCA2-deficient DLD1 cell line (Figure S5B and S5C). Moreover, inhibition of mutagenic TLS by treatment with JH-RE-06 showed additive cytotoxicity when combined with the depletion of EXO1 in the BRCA1-deficient cancer cell line (Figure 6B), thus further supporting the idea that EXO1 and TLS polymerases operate in distinct, complementary, pathways.

**Figure 6.**
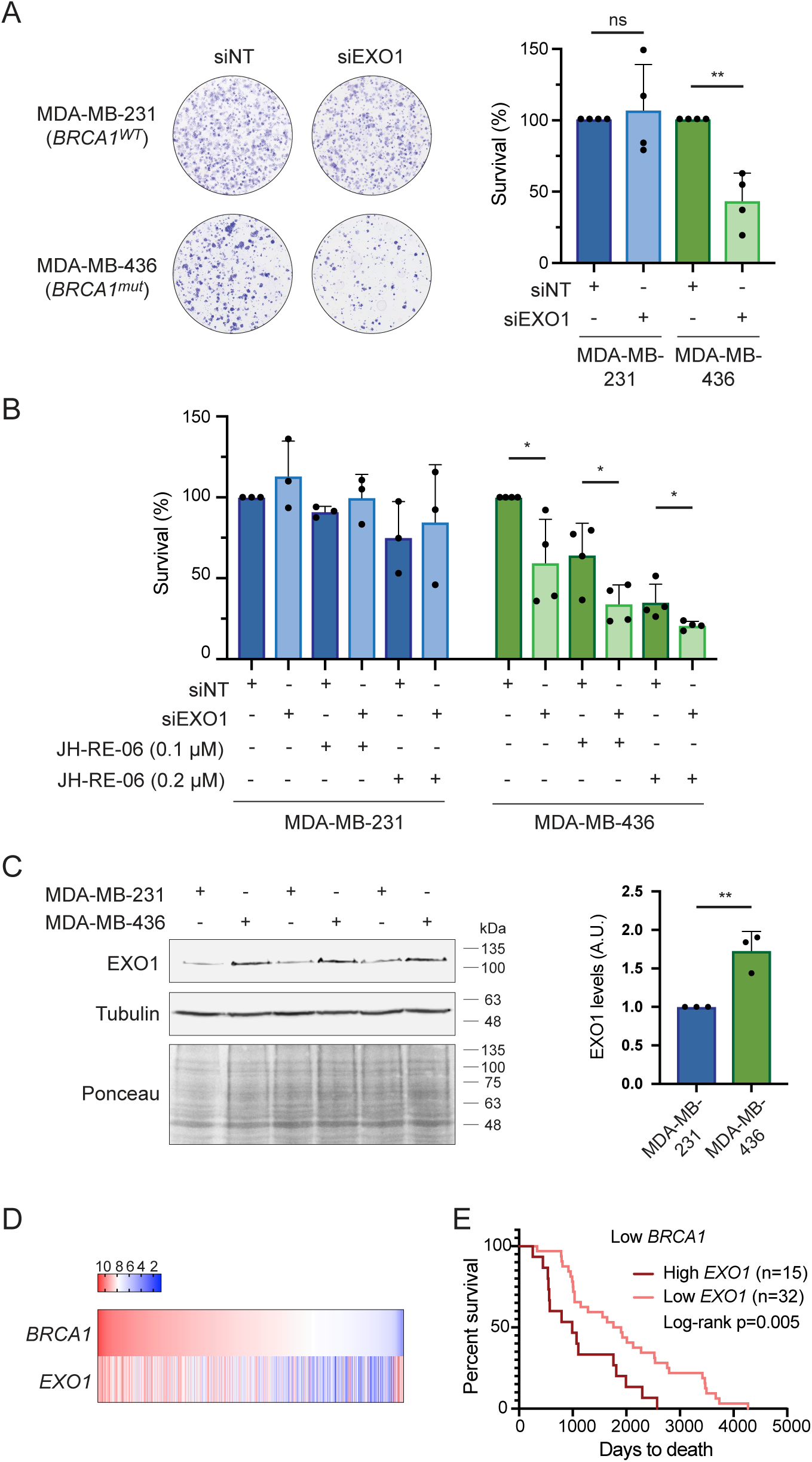
EXO1 is essential for BRCA1-mutant cells. **A.** Viability of MDA-MB-231 (*BRCA1^WT^*) and MDA-MB-436 (*BRCA1^mut^*) cells after transfection with the indicated siRNAs. Left: Representative images. Right: Quantification of cell survival as percentage of viable cells relative to siRNA control. Bars represent the means ± SD of independent biological replicates (n=4). Values of individual experiments are indicated as dots. p values were calculated using the Student’s *t*-test (ns: p>0.05; **: p<0.01). **B.** Viability of MDA-MB-231 and MDA-MB-436 cells after transfection with the indicated siRNAs upon treatment with JH-RE-06 at the indicated concentrations. Bars represent the means ± SD of independent biological replicates (n=3 for MDA-MB-231, n=4 for MDA-MB-436). Values of individual experiments are indicated as dots. p values were calculated using the Student’s *t*-test (ns: p>0.05; *: p<0.05). **C.** Western blot showing EXO1 protein levels in MDA-MB-231 and MDA-MB-436 cells. Tubulin and Ponceau served as loading controls. Quantifications of EXO1 levels, normalized to tubulin, are shown on the right. Bars represent the means ± SD of independent biological replicates (n=3). p value was calculated using the Student’s *t*-test (**: p<0.01). **D.** Heat map of the correlation between *BRCA1* and *EXO1* expressions retrieved from TCGA Breast Cancer (BRCA) dataset using UCSC Xena (n=1218). **E**. Survival of breast cancer patients with low expression levels of *BRCA1*, stratified according to their *EXO1* levels in two categories: high (dark red) or low (light red). Data from TCGA BRCA dataset. The number of samples is indicated.

EXO1 has been reported to be deregulated in several cancers (49). Indeed, data from The Cancer Genome Atlas (TCGA) shows that *EXO1* is overexpressed in breast cancer tissues compared to control (Figure S5D). Therefore, we hypothesized that BRCA1-deficient cells might be particularly addicted to *EXO1* overexpression. In fact, we observed a significant increase in EXO1 protein levels in BRCA1-deficient compared to BRCA1-proficient cells (Figure 6C). Furthermore, through a comparison of *EXO1* and *BRCA1* mRNA expression levels from TCGA breast cancer (BRCA) dataset, we observed that breast tumor samples exhibiting extremely low expression of *BRCA1* gene tended to display an elevated expression of *EXO1* (Figure 6D). However, this link was not as prominently observed in breast cancer patients with low expression of *BRCA2* (Figure S5E). Additionally, we observed that breast cancer patients with tumors expressing lower *BRCA1* levels had a more favorable prognosis specifically within the cohort exhibiting lower expression levels of *EXO1* (Figure 6E). Hence, our findings revealed that elevated *EXO1* expression correlates with BRCA1 deficiency and promotes tumor survival. These results open up the possibility of exploiting EXO1 inhibition in the clinic as a potential therapeutic strategy for BRCA1-deficient tumors. Notably, these findings do not appear to be exclusive to EXO1 but are shared with other factors involved in long-range resection since we also found elevated expression of *DNA2* and *BLM* in breast tumor samples with very low *BRCA1* expression, while this phenomenon was not as evident in the case of the short-range resection factor *MRE11* (Figure S6A). Again, this link was not observed in breast cancer patients with low expression of *BRCA2* (Figure S6B). Collectively, these results suggest that BRCA1-deficient cancer cells, in contrast to BRCA2-deficient cells, require a hyper-activation of long-range DNA resection factors for their survival.

## DISCUSSION

While it is widely accepted that RPA-coated ssDNA is the signal that activates the ATR checkpoint upon replication stress, the exact origin of this signal remains less understood. The prevailing model posits that ssDNA accumulates at replication forks as a result of a functional uncoupling of the MCM helicase from the replicative polymerases (50). However, this model does not consider the possibility of repriming. Here, we present data supporting an alternative model whereby the activation of the ATR checkpoint upon replicative stress occurs at post-replicative ssDNA gaps, expanded by EXO1 and DNA2, rather than directly at replication forks due to helicase uncoupling. Intriguingly, we observed only a limited contribution of human MRE11 on checkpoint activation, which contrasts with previous studies suggesting a role of this protein in gap expansion (16, 44). Thus, our observations indicate that the generation of sufficient ssDNA for ATR activation is primarily driven by factors involved in long-range resection, rather than MRE11-dependent short-range resection. However, whether EXO1 and/or DNA2 can compensate the absence of MRE11 remains to be explored. Notably, the proposed model of checkpoint activation is not limited to human cells but appears to be more general, as García-Rodríguez and colleagues previously identified processed-ssDNA gaps as the source of damage checkpoint signaling in *Saccharomyces cerevisiae* (24). Importantly, nuclease activity targeting ssDNA gaps must be carefully controlled to prevent the harmful consequences of over-resection. In this context, ATR activation has been demonstrated to initiate a negative feedback mechanism that prevents excessive resection via phosphorylation of, at least, EXO1 (24, 51).

The generation of these ssDNA gaps may depend on factors such as timing, the dosage of the genotoxic agent, or the genetic background (4). We demonstrated that, in our experimental setup, gaps were generated upon exposure to MMS without a noticeable formation of DSBs. We then established a link in *exo1* cells between the defective checkpoint activation and the extension of gap size, as we observed that DNA tracts became insensitive to S1 nuclease digestion following MMS treatment (Figure 5A), possibly due to ineffective digestion when gaps are shorter. As an alternative explanation, the use of DNA spreading may not facilitate the separation of sister chromatids, potentially hindering the detection of DNA tract shortening by S1 treatment unless there is an overlap of ssDNA regions in both adjacent sister chromatids, as shown in recent work from the Caldecott and Vindigni labs (52). Thus, one can hypothesize that, in the absence of EXO1, gaps are less likely to overlap and become undetectable by S1 treatment on DNA spreads. Remarkably, ssDNA gaps seem to form not only upon exposure to agents causing polymerase-blocking lesions but also in response to lesion-less replication stress induced by HU (4, 16, 40). It remains unclear why repriming is the chosen pathway for DDT in situations where replication forks do not face DNA lesions. However, repriming may result from the way PrimPol is recruited to long ssDNA stretches at stalled forks, which relies on its interaction with RPA (53, 54).

In addition, we found that proper ATR checkpoint activation upon MMS-induced replicative stress depends on the interaction between EXO1 and the DNA replication clamp PCNA. This interaction could facilitate the recruitment of EXO1 at sites of DNA damage and/or stimulate its processivity, mirroring observations seen in response to DSBs (33). Importantly, while PCNA is specifically loaded at the 3’-junction of the gap, EXO1 possess 5’-3’ exonuclease as well as 5’ structure-specific DNA endonuclease activities (55–57). Therefore, to accommodate the biochemical properties of both proteins, one might speculate that EXO1 could reach the 5’-junction of the gap if a ssDNA loop is formed, similar to those observed during the DNA end processing of DSBs, as well as in the context of base excision repair (58, 59). Notably, the ubiquitylation status of PCNA dictates DDT pathway choice: whereas monoubiquitylation of PCNA activates TLS, polyubiquitylation triggers TS through a mechanism that is still poorly understood (55). Interestingly, EXO1-dependent gap expansion is required to stimulate recombination events during TS (48, 60), raising the question whether polyubiquitylation of PCNA further enhances interaction with EXO1.

Consistent with a role in TS, we observed that *exo1* cells exhibited remarkable sensitive to the inhibition of the REV1-Polζ-dependent TLS. TLS polymerases are known to facilitate continuous DNA synthesis, thus limiting the accumulation of gaps induced by factors such as replication stress, oncogenes, or chemotherapy (14). Consequently, TLS inhibitors not only synergizes with gap-inducing conditions but also with other gap-filling pathways, highlighting their potential in cancer therapy since gaps have been identified as a cancer vulnerability (61). Given the association of EXO1 mutations with numerous cancer types (49), the inhibition of TLS could potentially offer therapeutic benefits in treating these cancers.

Strikingly, along with Kooij and colleagues (62), we revealed a novel synthetically lethal interaction between BRCA1 and EXO1. Recent studies have suggested that distinct BRCA1 functions prevent the toxic accumulation of ssDNA gaps. For instance, gaps have been proposed to arise in BRCA1-defiecient cells due to defects in Okazaki fragment processing (19). More recently, we collaborated in a study that found that BRCA1, when partnered with BARD1, facilitates continuous DNA synthesis in unchallenged conditions by directly ubiquitylating PCNA (30). Furthermore, BRCA1 initiates the repair of PrimPol-generated ssDNA gaps through TS and protects their integrity by limiting MRE11 activity, thus allowing the process of gap filling (26, 37). Yet, whether the shared role of EXO1 and BRCA1 in promoting gap filling via TS drives their synthetic lethality requires further investigation. Intriguingly, we discovered that the absence of EXO1 did not result in synthetic lethality in the context of BRCA2 deficiency, even though gaps also accumulate in BRCA2-deficient cells when exposed to replicative stress (25). This observation might be attributed to the fact that, while both BRCA1 and BRCA2 are crucial for HR, it is BRCA1, and not BRCA2, that facilitates resection (63). Remarkably, we found that EXO1 is upregulated in BRCA1-deficient cells, and gene expression levels of *EXO1*, *DNA2* and *BLM* are elevated in tumor samples with very low *BRCA1* expression. These observations suggest the activation of a compensatory mechanism involving long-range resection in BRCA1-mutant cells. Accordingly, BLM, which cooperates with the nuclease DNA2 during long-range resection, has also been found to be essential in BRCA1-deficient but not in BRCA1-proficient cells (62). Thus, we believe that BRCA1-mutant cells rely not only on TLS polymerases, given the reported synthetic lethality upon loss of REV1-Polζ or POLQ (14, 26–29), but also on proficient long-range resection for the repair of ssDNA gaps. Noteworthy, EXO1 might also play a role in facilitating the recruitment of TLS polymerases, as demonstrated for Polτ and Polκ at ssDNA gaps intermediates generated during nucleotide excision repair (64). In any case, the accumulation of unrepaired ssDNA gaps, together with impaired ATR checkpoint activation, can result in the exhaustion of the RPA pool. Such scenario has been shown to trigger DNA breakage and, consequently, cell death (65). Furthermore, in agreement with a role of ATR in protecting ssDNA gaps from EXO1 over-resection, ATR inhibition has been shown to exacerbate the PARPi-induction of ssDNA gaps in BRCA1-deficient cells (19, 66). Importantly, we showed an additive effect of the loss of EXO1 and the inhibition of the REV1-Polζ-dependent TLS in the survival of BRCA1-deficient cells. Thus, our findings suggests that BRCA1-mutant patients might benefit from combinational therapies involving inhibitors targeting TLS along with EXO1 inhibition, such as the recently described (67).

## DATA AVAILABILITY

The data underlying this article are available in the article and in its online supplementary material. Raw data will be shared on reasonable request to the corresponding authors.

## ACKNOWLEDGMENTS

We thank Lene J. Rasmussen for kindly providing the YFP-EXO1 plasmid.

## Author contributions

N. G-R.: Conceptualization, Investigation, Formal Analysis, Writing (original draft), Writing (review and editing), Funding Acquisition. MdC. D-P.: Investigation. P. H.: Writing (review and editing), Funding Acquisition, Supervision.

## FIGURE LEGENDS

**Figure S1. Loss of PrimPol sensitizes RPE1 cells to MMS. A**. PrimPol knockdown corresponding to the experiment shown in Figure 1C. Western blot for monitoring PrimPol levels 48h after transfection with control (siNT) or PrimPol siRNA. Tubulin served as loading control. B. Viability of RPE1 cells after transfection with the indicated siRNAs, upon treatment with 0.02 mM MMS, relative to control DMSO-treated cells. Bars represent the means ± SD of independent biological replicates (n=3). Values of individual experiments are indicated as dots. p values were calculated using the Student’s *t*-test (*: p<0.05).

**Figure S2. EXO1 function is required for robust ATR checkpoint activation upon replicative stress. A**. Western blots for monitoring CHK1-S345 phosphorylation in WT and *exo1* cells upon addition of 4-NQO (0.25 μM). Actin served as loading control. Quantifications of CHK1-pS345 intensities, normalized to actin, are shown on the middle. Bars represent the means ± SD of independent biological replicates (n=3). p-values were calculated using the Student’s *t*-test (*: p<0.05; **: p<0.01). **B**. Western blots for monitoring CHK1-S345 phosphorylation in WT and *exo1* cells upon addition of MMS (left, 2 mM) or HU (right, 2 mM) up to 3 hours. Tubulin served as loading control. Quantification of CHK1-S345 phosphorylation relative to tubulin is shown on the bottom. **C**. Western blots for monitoring CHK1-S345 phosphorylation in WT and *exo1* KO clones #15 and #9 upon addition of MMS (2 mM). Tubulin served as loading control. Cell cycle profiles are shown on the right. **D** and **E**. Western blots for monitoring CHK1-S345 phosphorylation in U2OS (C) or HeLa (D) cells treated with the indicated siRNAs upon addition of MMS (left, 2 mM) or HU (right, 2 mM). Actin served as loading control. Cell cycle profiles are shown on the right. **F**. Histogramas of CHK1-pS345 intensities in WT (red) and *exo1* (blue) cells in S-phase, gated from the flow cytometry experiment shown in Figure 2D.

Figure S3. **Treatments with MMS, HU, or 4-NQO do not lead to noticeable formation of DSBs. A**. Western blot for monitoring the levels of the indicated proteins upon addition of CPT (1 μM), MMS (2 mM), HU (2 mM) or 4-NQO (0.25 μM). Tubulin served as loading control. **B**. Analysis of the formation of 53BP1 foci upon the addition of the indicated genotoxic agents for 60 min. Scale bar, 25 μM.

**Figure S4. MRE11 is dispensable for ATR checkpoint activation upon replicative stress. A.** Western blots for monitoring CHK1-S345 phosphorylation in WT (RPE1) cells treated with control (siNT) or MRE11 siRNA, upon the addition of MMS (2 mM). Tubulin served as loading control. Quantifications of CHK1-pS345 intensities, normalized to tubulin, are shown at the right. Bars represent the means ± SD of independent biological replicates (n=3). p-values were calculated using the Student’s *t*-test (ns: p>0.05). **B.** Cell cycle profiles of siNT or siMRE11-treated cells. **C.** Western blots for monitoring CHK1-S345 phosphorylation in WT cells treated with control (DMSO) or mirin (25 μM), upon the addition of MMS (2 mM). Actin served as loading control.

**Figure S5. EXO1 loss is essential for BRCA1 but not BRCA2-mutant cells. A.** Western blot analysis of the lysates of the MDA-MB-231 and MDA-MB-436 cells studied in Figure 6A. **B**. Viability of DLD-1 and DLD-1 *BRCA2^-/-^* cells after transfection with the indicated siRNAs. Left: Representative images. Right: Quantification of cell survival as percentage of viable cells relative to siRNA control.. Bars represent the means ± SD of independent biological replicates (n=3). Values of individual experiments are indicated as dots. p values were calculated using the Student’s *t*-test (ns: p>0.05). **C**. Western blot analysis of the lysates of the DLD-1 and DLD-1 *BRCA2^-/-^* cells studied in Figure S4B. **D**. *EXO1* mRNA expression levels in breast cancer samples (n=1085, red) compared to non-tumor samples (n=291, grey), retrieved from TCGA and GTEx datasets and analyzed by GEPIA platform. **E**. Heat map of the correlation between *BRCA2* and *EXO1* expressions retrieved from TCGA Breast Cancer (BRCA) dataset using UCSC Xena (n=1218).

**Figure S6. Low *BRCA1* expression correlates with elevated expression levels of the long-range resection factors *DNA2* and *BLM* in breast tumor samples.** Heat map of the correlation between the expressions of *BRCA1*, *DNA2*, *BLM* and *MRE11* (**A**), as well as the correlation between the expressions of *BRCA2*, *DNA2*, *BLM* and *MRE11* (**B**). Data retrieved from TCGA Breast Cancer (BRCA) dataset using UCSC Xena (n=1218).

## REFERENCES

1. Zeman, M.K. and Cimprich, K.A. (2014) Causes and consequences of replication stress. Nature Cell Biology, 16, 2–9.

2. Zou, L. and Elledge, S.J. (2003) Sensing DNA Damage Through ATRIP Recognition of RPA-ssDNA Complexes. Science, 300, 1542–1548.

3. Maréchal, A. and Zou, L. (2013) DNA damage sensing by the ATM and ATR kinases. Csh Perspect Biol, 5, a012716–a012716.

4. Quinet, A., Tirman, S., Cybulla, E., Meroni, A. and Vindigni, A. (2021) To skip or not to skip: choosing repriming to tolerate DNA damage. Mol Cell, 10.1016/j.molcel.2021.01.012.

5. Friedberg, E.C. (2005) Suffering in silence: the tolerance of DNA damage. Nature Reviews Molecular Cell Biology, 6, 943–953.

6. Sale, J.E. (2013) Translesion DNA Synthesis and Mutagenesis in Eukaryotes. Cold Spring Harb. Perspect. Biol., 5, a012708.

7. Vaisman, A. and Woodgate, R. (2017) Translesion DNA polymerases in eukaryotes: what makes them tick? Crit. Rev. Biochem. Mol. Biol., 52, 274–303.

8. Guilliam, T.A. and Doherty, A.J. (2017) PrimPol-Prime Time to Reprime. Genes, 8.

9. Bianchi, J., Rudd, S.G., Jozwiakowski, S.K., Bailey, L.J., Soura, V., Taylor, E., Stevanovic, I., Green, A.J., Stracker, T.H., Lindsay, H.D., et al. (2013) PrimPol bypasses UV photoproducts during eukaryotic chromosomal DNA replication. Mol Cell, 52, 566–73.

10. García-Gómez, S., Reyes, A., Martínez-Jiménez, M.I., Chocrón, E.S., Mourón, S., Terrados, G., Powell, C., Salido, E., Méndez, J., Holt, I.J., et al. (2013) PrimPol, an archaic primase/polymerase operating in human cells. Mol Cell, 52, 541–53.

11. Mourón, S., Rodriguez-Acebes, S., Martínez-Jiménez, M.I., García-Gómez, S., Chocrón, S., Blanco, L. and Méndez, J. (2013) Repriming of DNA synthesis at stalled replication forks by human PrimPol. Nature Structural & Molecular Biology, 20, 1383–1389.

12. Wong, R.P., García-Rodríguez, N., Zilio, N., Hanulová, M. and Ulrich, H.D. (2020) Processing of DNA Polymerase-Blocking Lesions during Genome Replication Is Spatially and Temporally Segregated from Replication Forks. Mol Cell, 77, 3–16.e4.

13. Zellweger, R., Dalcher, D., Mutreja, K., Berti, M., Schmid, J.A., Herrador, R., Vindigni, A. and Lopes, M. (2015) Rad51-mediated replication fork reversal is a global response to genotoxic treatments in human cells. The Journal of Cell Biology, 208, 563–579.

14. Nayak, S., Calvo, J.A., Cong, K., Peng, M., Berthiaume, E., Jackson, J., Zaino, A.M., Vindigni, A., Hadden, M.K. and Cantor, S.B. (2020) Inhibition of the translesion synthesis polymerase REV1 exploits replication gaps as a cancer vulnerability. Sci Adv, 6, eaaz7808.

15. Elvers, I., Johansson, F., Groth, P., Erixon, K. and Helleday, T. (2011) UV stalled replication forks restart by re-priming in human fibroblasts. Nucleic Acids Research, 39, 7049–7057.

16. Piberger, A.L., Bowry, A., Kelly, R.D.W., Walker, A.K., González-Acosta, D., Bailey, L.J., Doherty, A.J., Méndez, J., Morris, J.R., Bryant, H.E., et al. (2020) PrimPol-dependent single-stranded gap formation mediates homologous recombination at bulky DNA adducts. Nat Commun, 11, 5863.

17. Quinet, A., Tirman, S., Jackson, J., Šviković, S., Lemaçon, D., Carvajal-Maldonado, D., González-Acosta, D., Vessoni, A.T., Cybulla, E., Wood, M., et al. (2020) PRIMPOL-Mediated Adaptive Response Suppresses Replication Fork Reversal in BRCA-Deficient Cells. Mol. Cell, 77, 461–474.e9.

18. González-Acosta, D., Blanco-Romero, E., Ubieto-Capella, P., Mutreja, K., Míguez, S., Llanos, S., García, F., Muñoz, J., Blanco, L., Lopes, M., et al. (2021) PrimPol-mediated repriming facilitates replication traverse of DNA interstrand crosslinks. Embo J, 10.15252/embj.2020106355.

19. Cong, K., Peng, M., Kousholt, A.N., Lee, W.T.C., Lee, S., Nayak, S., Krais, J., VanderVere-Carozza, P.S., Pawelczak, K.S., Calvo, J., et al. (2021) Replication gaps are a key determinant of PARP inhibitor synthetic lethality with BRCA deficiency. Mol Cell, 10.1016/j.molcel.2021.06.011.

20. Bai, G., Kermi, C., Stoy, H., Schiltz, C.J., Bacal, J., Zaino, A.M., Hadden, M.K., Eichman, B.F., Lopes, M. and Cimprich, K.A. (2020) HLTF Promotes Fork Reversal, Limiting Replication Stress Resistance and Preventing Multiple Mechanisms of Unrestrained DNA Synthesis. Mol Cell, 78, 1237–1251.e7.

21. Šviković, S., Crisp, A., Tan-Wong, S.M., Guilliam, T.A., Doherty, A.J., Proudfoot, N.J., Guilbaud, G. and Sale, J.E. (2019) R-loop formation during S phase is restricted by PrimPol-mediated repriming. Embo J, 38.

22. Tercero, J., Longhese, M. and Diffley, J. (2003) A Central Role for DNA Replication Forks in Checkpoint Activation and Response. Molecular Cell, 11, 1323–1336.

23. Shimada, K., Pasero, P. and Gasser, S.M. (2002) ORC and the intra-S-phase checkpoint: a threshold regulates Rad53p activation in S phase. Genes & development, 16, 3236–52.

24. García-Rodríguez, N., Morawska, M., Wong, R.P., Daigaku, Y. and Ulrich, H.D. (2018) Spatial separation between replisome- and template-induced replication stress signaling. The EMBO journal, 37.

25. Panzarino, N.J., Krais, J.J., Cong, K., Peng, M., Mosqueda, M., Nayak, S.U., Bond, S.M., Calvo, J.A., Doshi, M.B., Bere, M., et al. (2021) Replication Gaps Underlie BRCA Deficiency and Therapy Response. Cancer Res, 81, 1388–1397.

26. Taglialatela, A., Leuzzi, G., Sannino, V., Cuella-Martin, R., Huang, J.-W., Wu-Baer, F., Baer, R., Costanzo, V. and Ciccia, A. (2021) REV1-Polζ maintains the viability of homologous recombination-deficient cancer cells through mutagenic repair of PRIMPOL-dependent ssDNA gaps. Mol Cell, 10.1016/j.molcel.2021.08.016.

27. Belan, O., Sebald, M., Adamowicz, M., Anand, R., Vancevska, A., Neves, J., Grinkevich, V., Hewitt, G., Segura-Bayona, S., Bellelli, R., et al. (2022) POLQ seals post-replicative ssDNA gaps to maintain genome stability in BRCA-deficient cancer cells. Mol Cell, 10.1016/j.molcel.2022.11.008.

28. Schrempf, A., Bernardo, S., Verge, E.A.A., Otero, M.A.R., Wilson, J., Kirchhofer, D., Timelthaler, G., Ambros, A.M., Kaya, A., Wieder, M., et al. (2022) POLθ processes ssDNA gaps and promotes replication fork progression in BRCA1-deficient cells. Cell Reports, 10.1016/j.celrep.2022.111716.

29. Mann, A., Ramirez-Otero, M.A., Antoni, A.D., Hanthi, Y.W., Sannino, V., Baldi, G., Falbo, L., Schrempf, A., Bernardo, S., Loizou, J., et al. (2022) POLθ prevents MRE11-NBS1-CtIP-dependent fork breakage in the absence of BRCA2/RAD51 by filling lagging-strand gaps. Mol Cell, 82, 4218–4231.e8.

30. Salas-Lloret, D., García-Rodríguez, N., Giebel, L., Ru, A. de, Veelen, P.A. van, Huertas, P., Vertegaal, A.C.O. and González-Prieto, R. (2023) BRCA1/BARD1 ubiquitinates PCNA in unperturbed conditions to promote replication fork stability and continuous DNA synthesis. bioRxiv, 10.1101/2023.01.12.523782.

31. Cantor, S.B. (2021) Revisiting the BRCA-pathway through the lens of replication gap suppression “Gaps determine therapy response in BRCA mutant cancer.” Dna Repair, 107, 103209.

32. Liberti, S.E., Andersen, S.D., Wang, J., May, A., Miron, S., Perderiset, M., Keijzers, G., Nielsen, F.C., Charbonnier, J.-B.B., Bohr, V.A., et al. (2011) Bi-directional routing of DNA mismatch repair protein human exonuclease 1 to replication foci and DNA double strand breaks. DNA repair, 10, 73–86.

33. Chen, X., Paudyal, S.C., Chin, R.-I. and You, Z. (2013) PCNA promotes processive DNA end resection by Exo1. Nucleic Acids Research, 41, 9325–9338.

34. Guzmán, C., Bagga, M., Kaur, A., Westermarck, J. and Abankwa, D. (2014) ColonyArea: An ImageJ Plugin to Automatically Quantify Colony Formation in Clonogenic Assays. Plos One, 9, e92444.

35. Goldman, M.J., Craft, B., Hastie, M., Repečka, K., McDade, F., Kamath, A., Banerjee, A., Luo, Y., Rogers, D., Brooks, A.N., et al. (2020) Visualizing and interpreting cancer genomics data via the Xena platform. Nat. Biotechnol., 38, 675– 678.

36. Tang, Z., Li, C., Kang, B., Gao, G., Li, C. and Zhang, Z. (2017) GEPIA: a web server for cancer and normal gene expression profiling and interactive analyses. Nucleic Acids Res, 45, W98–W102.

37. Tirman, S., Quinet, A., Wood, M., Meroni, A., Cybulla, E., Jackson, J., Pegoraro, S., Simoneau, A., Zou, L. and Vindigni, A. (2021) Temporally distinct post-replicative repair mechanisms fill PRIMPOL-dependent ssDNA gaps in human cells. Mol Cell, 81, 4026–4040.e8.

38. Quinet, A., Carvajal-Maldonado, D., Lemacon, D. and Vindigni, A. (2017) Chapter Three DNA Fiber Analysis: Mind the Gap! Methods Enzymol, 591, 55–82.

39. Kobayashi, K., Guilliam, T.A., Tsuda, M., Yamamoto, J., Bailey, L.J., Iwai, S., Takeda, S., Doherty, A.J. and Hirota, K. (2016) Repriming by PrimPol is critical for DNA replication restart downstream of lesions and chain-terminating nucleosides. Cell Cycle, 15, 1–12.

40. Gallo, D., Kim, T., Szakal, B., Saayman, X., Narula, A., Park, Y., Branzei, D., Zhang, Z. and Brown, G.W. (2019) Rad5 Recruits Error-Prone DNA Polymerases for Mutagenic Repair of ssDNA Gaps on Undamaged Templates. Molecular cell, 10.1016/j.molcel.2019.01.001.

41. Huertas, P. (2010) DNA resection in eukaryotes: deciding how to fix the break. Nature Structural & Molecular Biology, 17, 11–6.

42. Hsiang, Y.H., Lihou, M.G. and Liu, L.F. (1989) Arrest of replication forks by drug-stabilized topoisomerase I-DNA cleavable complexes as a mechanism of cell killing by camptothecin. Cancer Res., 49, 5077–82.

43. Cejka, P. (2015) DNA End Resection: Nucleases Team Up with the Right Partners to Initiate Homologous Recombination*. J. Biological Chem., 290, 22931–22938.

44. Hashimoto, Y., Chaudhuri, A.R., Lopes, M. and Costanzo, V. (2010) Rad51 protects nascent DNA from Mre11-dependent degradation and promotes continuous DNA synthesis. Nat. Struct. Mol. Biology, 17, 1305–1311.

45. Liu, W., Zhou, M., Li, Z., Li, H., Polaczek, P., Dai, H., Wu, Q., Liu, C., Karanja, K.K., Popuri, V., et al. (2016) A Selective Small Molecule DNA2 Inhibitor for Sensitization of Human Cancer Cells to Chemotherapy. EBioMedicine, 6, 73–86.

46. García-Rodríguez, N., Wong, R.P. and Ulrich, H.D. (2018) The helicase Pif1 functions in the template switching pathway of DNA damage bypass. Nucleic Acids Research, 46, 8347–8356.

47. Karras, G., Fumasoni, M., Sienski, G., Vanoli, F., Branzei, D. and Jentsch, S. (2013) Noncanonical Role of the 9-1-1 Clamp in the Error-Free DNA Damage Tolerance Pathway. Molecular Cell, 49, 536–546.

48. Vanoli, F., Fumasoni, M., Szakal, B., Maloisel, L. and Branzei, D. (2010) Replication and recombination factors contributing to recombination-dependent bypass of DNA lesions by template switch. PLoS genetics, 6, e1001205.

49. Keijzers, G., Bakula, D., Petr, M.A., Madsen, N.G., Teklu, A., Mkrtchyan, G., Osborne, B. and Scheibye-Knudsen, M. (2018) Human Exonuclease 1 (EXO1) Regulatory Functions in DNA Replication with Putative Roles in Cancer. International journal of molecular sciences, 20.

50. Byun, T.S., Pacek, M., Yee, M., Walter, J.C. and Cimprich, K.A. (2005) Functional uncoupling of MCM helicase and DNA polymerase activities activates the ATR-dependent checkpoint. Genes & Development, 19, 1040–52.

51. Tomimatsu, N., Mukherjee, B., Harris, J.L., Boffo, F.L., Hardebeck, M.C., Potts, P.R., Khanna, K.K. and Burma, S. (2017) DNA-damage-induced degradation of EXO1 exonuclease limits DNA end resection to ensure accurate DNA repair. The Journal of biological chemistry, 292, 10779–10790.

52. Meroni, A., Wells, S.E., Fonseca, C., Chaudhuri, A.R., Caldecott, K.W. and Vindigni, A. (2023) DNA Combing versus DNA Spreading and the Separation of Sister Chromatids. bioRxiv Prepr. Serv. biology, 10.1101/2023.05.02.539129.

53. Guilliam, T.A., Brissett, N.C., Ehlinger, A., Keen, B.A., Kolesar, P., Taylor, E.M., Bailey, L.J., Lindsay, H.D., Chazin, W.J. and Doherty, A.J. (2017) Molecular basis for PrimPol recruitment to replication forks by RPA. Nat. Commun., 8, 15222.

54. Wan, L., Lou, J., Xia, Y., Su, B., Liu, T., Cui, J., Sun, Y., Lou, H. and Huang, J. (2013) hPrimpol1/CCDC111 is a human DNA primase-polymerase required for the maintenance of genome integrity. EMBO reports, 14, 1104–12.

55. Moldovan, G.-L.L., Pfander, B. and Jentsch, S. (2007) PCNA, the maestro of the replication fork. Cell, 129, 665–79.

56. Wilson, D.M., Coleman, M.A., Adamson, A.W., Christensen, M., Lamerdin, J.E. and Carney, J.P. (1998) Hex1: a new human Rad2 nuclease family member with homology to yeast exonuclease 1. Nucleic Acids Res., 26, 3762–3768.

57. Lee, B.-I. and Wilson, D.M. (1999) The RAD2 Domain of Human Exonuclease 1 Exhibits 5′ to 3′ Exonuclease and Flap Structure-specific Endonuclease Activities*. J. Biol. Chem., 274, 37763–37769.

58. Yoo, J., Lee, D., Im, H., Ji, S., Oh, S., Shin, M., Park, D. and Lee, G. (2021) The mechanism of gap creation by a multifunctional nuclease during base excision repair. Sci. Adv., 7, eabg0076.

59. Xue, C., Salunkhe, S.J., Tomimatsu, N., Kawale, A.S., Kwon, Y., Burma, S., Sung, P. and Greene, E.C. (2022) Bloom helicase mediates formation of large single–stranded DNA loops during DNA end processing. Nat. Commun., 13, 2248.

60. Piberger, A., Walker, A.K., Morris, J.R., Bryant, H.E. and Petermann, E. (2019) PrimPol-dependent single-stranded gap formation mediates homologous recombination at bulky DNA adducts. Biorxiv, 10.1101/773242.

61. Patel, S.M., Dash, R.C. and Hadden, M.K. (2021) Translesion synthesis inhibitors as a new class of cancer chemotherapeutics. Expert Opin. Investig. Drugs, 30, 13–24.

62. Kooij, B. van de, Schreuder, A., Pavani, R.S., Garzero, V., Hoeck, A.V., Alonso, M.S.M., Koerse, D., Wendel, T.J., Callen, E., Boom, J., et al. (2023) EXO1-mediated DNA repair by single-strand annealing is essential for BRCA1-deficient cells. Biorxiv, 10.1101/2023.02.24.529205.

63. Prakash, R., Zhang, Y., Feng, W. and Jasin, M. (2015) Homologous Recombination and Human Health: The Roles of BRCA1, BRCA2, and Associated Proteins. Cold Spring Harb. Perspect. Biol., 7, a016600.

64. Sertic, S., Mollica, A., Campus, I., Roma, S., Tumini, E., Aguilera, A. and Muzi-Falconi, M. (2018) Coordinated Activity of Y Family TLS Polymerases and EXO1 Protects Non-S Phase Cells from UV-Induced Cytotoxic Lesions. Molecular cell, 70, 34–47.e4.

65. Toledo, L., Altmeyer, M., Rask, M.-B., Lukas, C., Larsen, D., Povlsen, L., Bekker-Jensen, S., Mailand, N., Bartek, J. and Lukas, J. (2014) ATR Prohibits Replication Catastrophe by Preventing Global Exhaustion of RPA. Cell, 156, 374.

66. Leung, W., Simoneau, A., Saxena, S., Jackson, J., Patel, P.S., Limbu, M., Vindigni, A. and Zou, L. (2023) ATR protects ongoing and newly assembled DNA replication forks through distinct mechanisms. Cell Rep., 42, 112792.

67. Paiano, J., Zolnerowich, N., Wu, W., Pavani, R., Wang, C., Li, H., Zheng, L., Shen, B., Sleckman, B.P., Chen, B.-R., et al. (2021) Role of 53BP1 in end protection and DNA synthesis at DNA breaks. Genes Dev., 35, 1356–1367.

