## Supplementary material for "EXO1-mediated ssDNA gap expansion is essential for ATR activation and to maintain viability in BRCA1-deficient cells": Figures S1-S6

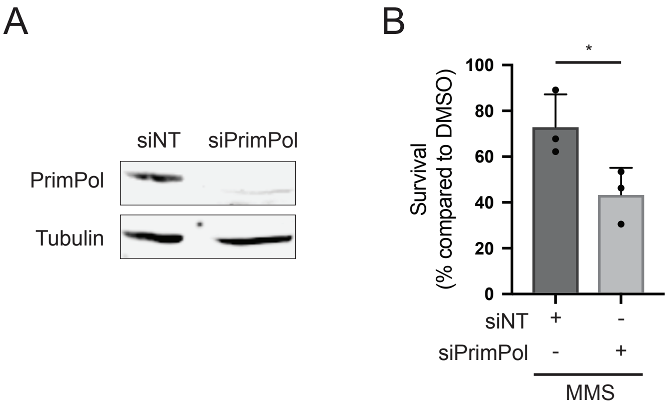


**Figure S1**. **Loss of PrimPol sensitizes RPE1 cells to MMS. A.** PrimPol knockdown corresponding to the experiment shown in Figure 1C. Western blots for monitoring PrimPol levels 48h after transfection with control (siNT) or PrimPol siRNA. Tubulin served as loading control. **B.** Viability of RPE1 cells after transfection with the indicated siRNAs upon treatment with 0.02 mM MMS, relative to control DMSO-treated cells. Bars respresnt the means ± SD of independent biological replicates (n=3). Values of individual experiments are indicated as dots. p values were calculated using the Student's *t*-test (*: $p<0.05$).

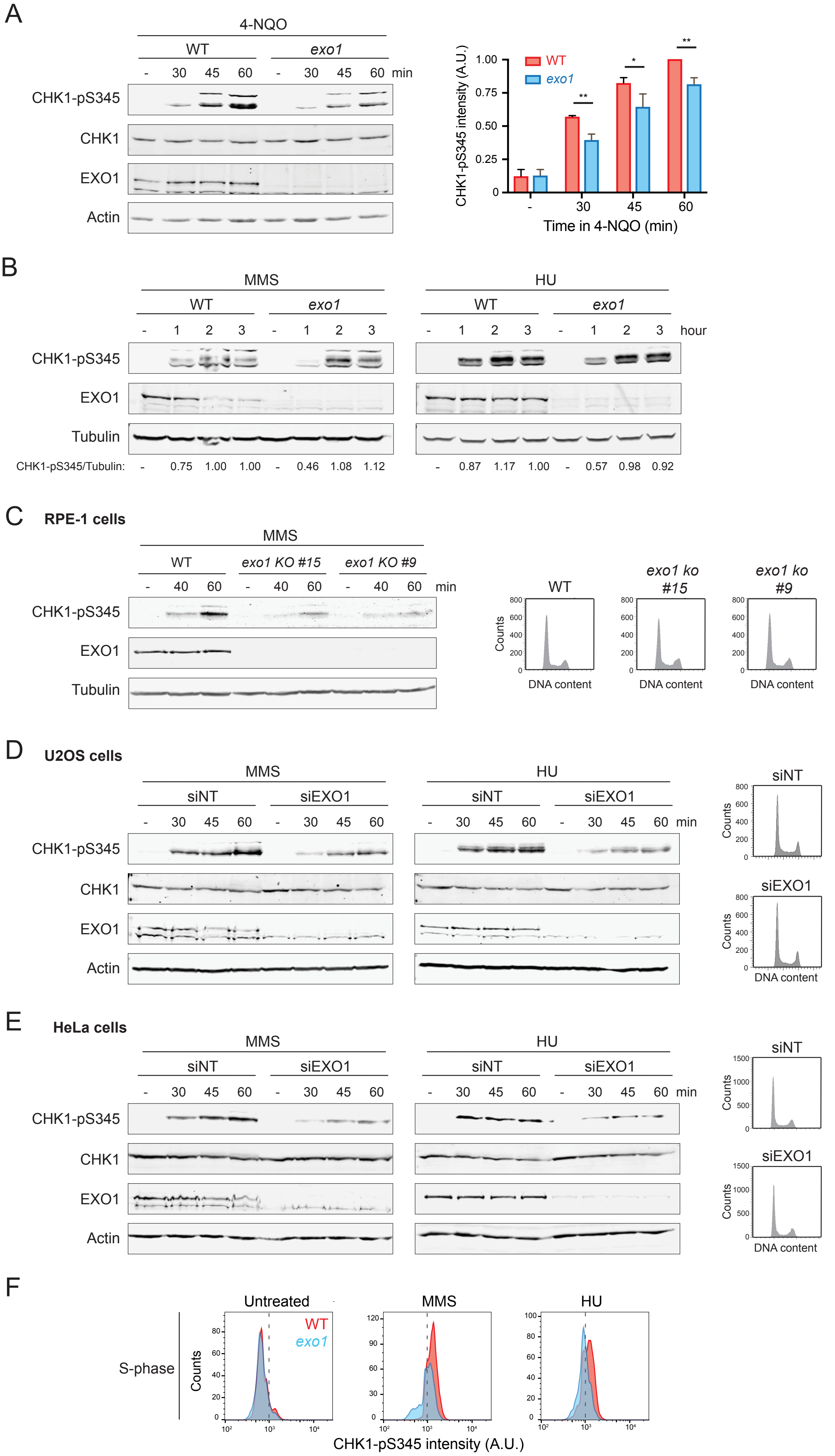
A
4-NQO
WT
exo1
- 30 45 60 - 30 45 60 min
CHK1-pS345
CHK1
EXO1
Actin
WT
exo1
CHK1-pS345 intensity (A.U.)
1.00
0.75
0.50
0.25
0
**
*
**
- 30 45 60
Time in 4-NQO (min)
B
MMS
HU
WT
exo1
- 1 2 3 - 1 2 3 - 1 2 3 - 1 2 3 hour
CHK1-pS345
EXO1
Tubulin
CHK1-pS345/Tubulin: - 0.75 1.00 1.00 - 0.46 1.08 1.12 - 0.87 1.17 1.00 - 0.57 0.98 0.92
C
RPE-1 cells
MMS
WT
exo1 KO #15
exo1 KO #9
- 40 60 - 40 60 - 40 60 min
CHK1-pS345
EXO1
Tubulin
WT
exo1 ko #15
exo1 ko #9
Counts
DNA content
D
U2OS cells
MMS
HU
siNT
siEXO1
- 30 45 60 - 30 45 60 - 30 45 60 - 30 45 60 min
CHK1-pS345
CHK1
EXO1
Actin
siNT
siEXO1
Counts
DNA content
E
HeLa cells
MMS
HU
siNT
siEXO1
- 30 45 60 - 30 45 60 - 30 45 60 - 30 45 60 min
CHK1-pS345
CHK1
EXO1
Actin
siNT
siEXO1
Counts
DNA content
F
Untreated
MMS
HU
WT
exo1
S-phase
Counts
CHK1-pS345 intensity (A.U.)

**Figure S2**. **EXO1 function is required for robust ATR checkpoint activation upon replicative stress. A.** Western blots for monitoring CHK1-S345 phosphorylation in WT and *exo1* ko cells upon addition of 4-NQO (0.25 μM). Actin served as loading control. Quantifications of CHK1-pS345 intensities, normalized to actin, are shown on the middle. Bars respresnt the means ± SD of independent biological replicates (n=3). p values were calculated using the Student's *t*-test (*: $p<0.05$; **: $p<0.01$). Cell cycle profiles are shown at the right. **B.** Western blots for monitoring CHK1-S345 phosphorylation in WT and *exo1* KO clones #15 and #9 upon addition of MMS (2mM). Tubulin served as loading control. Cell cycle profiles are shown onthe right. **C** and **D.** Western blots for monitoring CHK1-S345 phosphorylation in U2OS (C) or HeLa (D) cells treated with the indicated siRNAs upon addition of MMS (left, 2mM) or HU (right, 2mM). Actin served as loading control. Cell cycle profiles are shown at the right. **E.** Histograms of CHK1-pS345 intensities in WT (red) and *exo1* (blue) cells in S-phase, gated from the flow citometry experiment shown in Figure 2D.

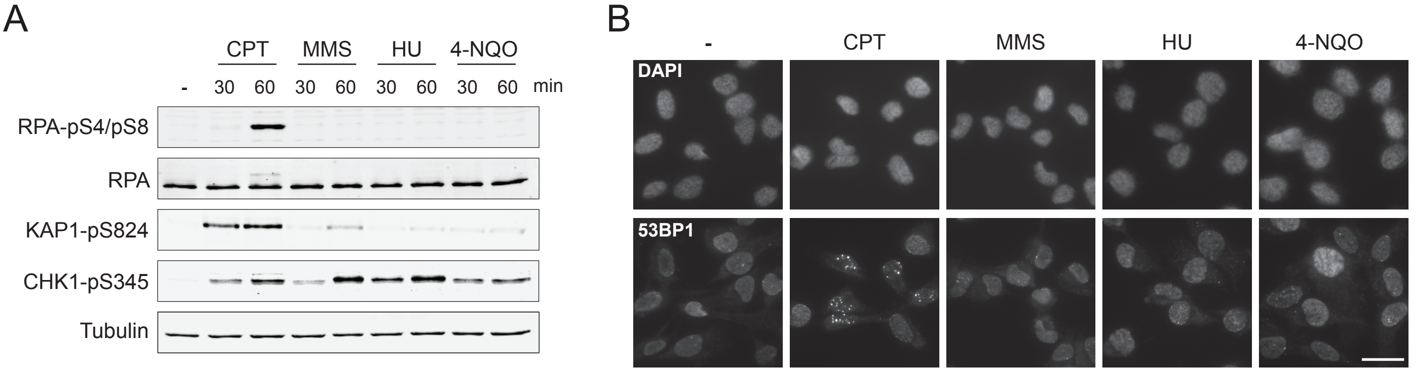


**Figure S3**. **Treatments with MMS, HU or 4-NQO do not lead to noticeable formation of DSBs. A.** Western blot for monitoring the levels of the indicated proteins upon addition of CPT (1 μM), MMS (2 mM), HU (2 mM) or 4-NQO (0.25 μM). Tubulin served as loading control. **B.** Analysis of the formation of 53BP1 foci upon the addition of the indicated genotoxic agents for 60 min. Scale bar, 25 μM.

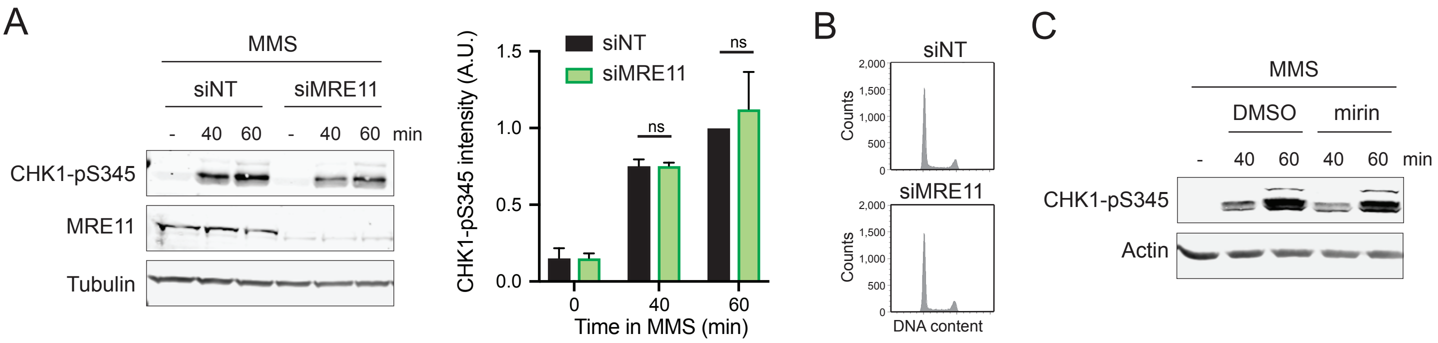


**Figure S4**. **MRE11 is dispensable for ATR checkpoint activation upon replicative stress. A.** Western blots for monitoring CHK1-S345 phosphorylation in WT (RPE1) cells treated with control (siNT) or MRE11 siRNA, upon the addition of MMS (2 mM). Tubulin served as loading control. Quantifications of CHK1-pS345 intensities, normalized to tubulin, are shown at the right. Bars represent the means ± SD of independent biological replicates (n=3). p values were calculated using the Student's *t*-test (ns: p>0.05). **B.** Cell cycle profiles of siNT or siMRE11-treated cells. **C.** Western blots for monitoring CHK1-S345 phosphorylation in WT cells treated with control (DMSO) or mirin (25 μM), upon the addition of MMS (2 mM). Actin served as loading control.

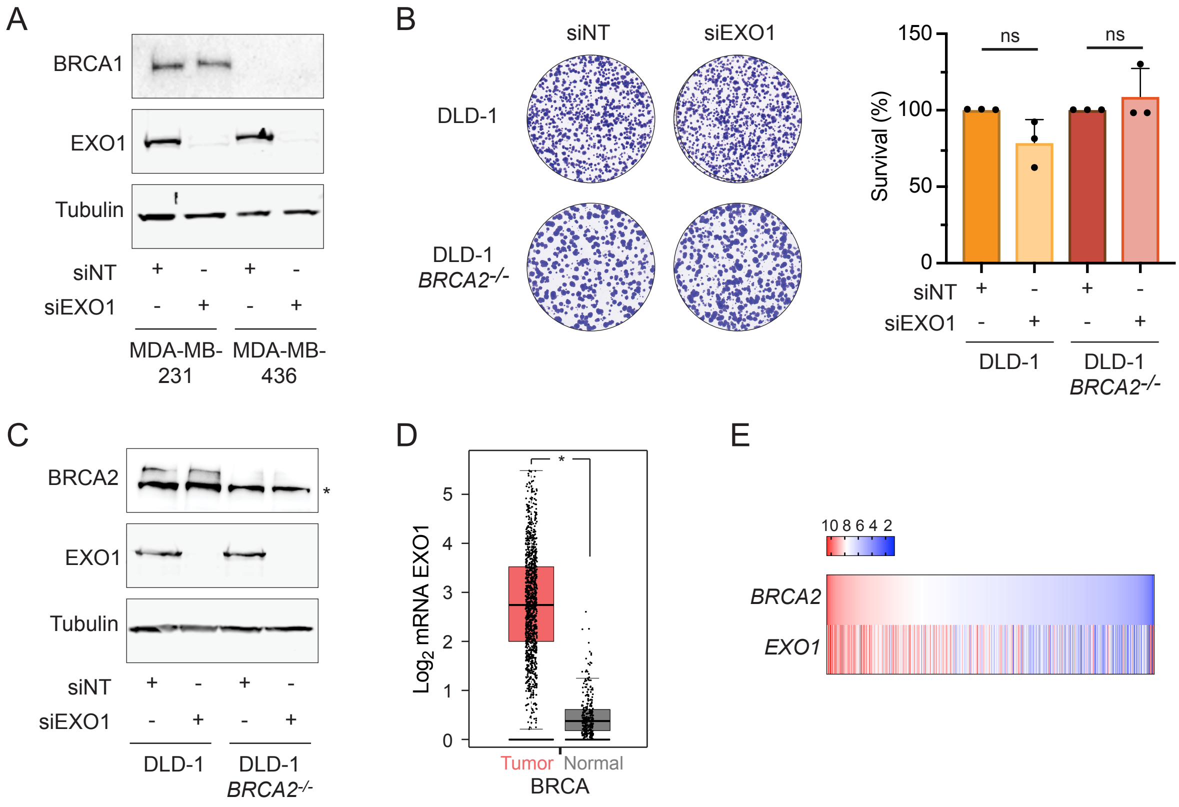


**Figure S5. EXO1 loss is essential for BRCA1 but not BRCA2-mutant cells. A.** Western blot analysis of the lysates of the MDA-MB-231 and MDA-MB-436 cells studied in Figure 6A. **B.** Viability of DLD-1 and DLD-1 *BRCA2-/-* cells after transfection with the indicated siRNAs. Left: Representative images. Right: Quantification of cell survival as percentage of viable cells relative to siRNA control. Bars respresnt the means ± SD of independent biological replicates (n=3). Values of individual experiments are indicated as dots. p values were calculated using the Student's *t*-test (ns: p>0.05). **C**.Western blot analysis of the lysates of the DLD-1 and DLD-1 *BRCA2-/-* cells studied in Figure S4B. **D**. *EXO1* mRNA expression levels in breast cancer samples (n=1085, red) compared to non-tumor samples (n= 291, grey), retrieved from TCGA and GTEx datasets and analyzed by GEPIA platform. **E**. Heat map of the correlation between *BRCA2* and *EXO1* expressions retrieved from TCGA Breast Cancer (BRCA) dataset using UCSC Xena (n=1218).

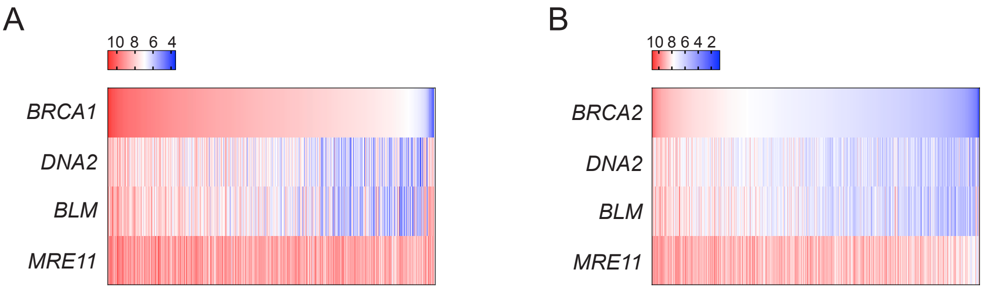


**Figure S6. Low *BRCA1* expression correlates with elevated expression levels of the long-range resection factors *DNA2 and BLM* in breast tumor samples.** Heat map of the correlation between the expressions of *BRCA1*, *DNA2, BLM and MRE11* (A), as well as the correlation between the expressions of *BRCA2*, *DNA2, BLM and MRE11* (B). Data retrieved from TCGA Breast Cancer (BRCA) dataset using UCSC Xena (n=1218).
